# Spatial Transcriptomics in Breast Cancer Reveals Tumour Microenvironment-Driven Drug Responses and Clonal Therapeutic Heterogeneity

**DOI:** 10.1101/2024.02.18.580660

**Authors:** María José Jiménez-Santos, Santiago García-Martín, Marcos Rubio-Fernández, Gonzalo Gómez-López, Fátima Al-Shahrour

## Abstract

Breast cancer is a heterogeneous disease that has the highest incidence and mortality rate among cancers in women worldwide. Breast cancer patients are stratified into three clinical subtypes with different treatment strategies and prognostic values. The development of targeted therapies against the biomarkers that define these strata constitutes one of the precedents of precision oncology, which aims to provide tailored treatments to cancer patients by targeting the molecular alterations found in each tumour. Although this approach has increased patient outcomes, many treatment failure cases still exist. Drug ineffectiveness and relapse have been associated with the coexistence of several malignant subpopulations with different drug sensitivities within the same lesion, a phenomenon known as intratumor heterogeneity. This heterogeneity has been extensively studied from a tumour-centric view, but recently, it has become evident that the tumour microenvironment plays a crucial role in intratumor heterogeneity. However, few studies consider the tumour-microenvironment interplay and its influence on drug sensitivity. In this work, we predict the sensitivity of 10x Visium spatial transcriptomics data from 9 breast cancer patients to >1,200 drugs and verify different response patterns across the tumour, interphase and microenvironment regions. We uncover a sensitivity continuum from the tumour core to the periphery accompanied by a functional gradient. Moreover, we identify conserved therapeutic clusters with distinct response patterns within the tumour region. We link the specific drug sensitivities of each therapeutic cluster to different ligand-receptor interactions that underpin distinct biological functions. Finally, we demonstrate that genetically identical cancer spots may belong to different therapeutic clusters and that this therapeutic heterogeneity is related to their location at the edge or core of tumour ducts. These results highlight the importance of considering the distance to the tumour core and the microenvironment composition when identifying suitable treatments to target intratumor heterogeneity.

## Background

Cancer is a group of genetic diseases that has been classically approached with a one-size-fits-all strategy, which often resulted in treatment inefficacy and adverse drug reactions [1], underlining the existence of intertumour heterogeneity [2]. Personalised precision oncology is a novel therapeutic strategy that aims to provide tailored treatments to cancer patients according to the molecular characteristics of individual tumours [3,4]. Breast cancer research has contributed to the foundations of personalised precision oncology by identifying biomarkers to guide patient stratification and selection of targeted therapies. In the clinical setting, an immunohistochemical analysis determines the expression of the oestrogen receptor (ER) and the progesterone receptor (PR) and the overexpression or amplification of the human epidermal growth factor receptor 2 (HER2). The status of these three hormone receptors guides the stratification of breast cancer patients into three groups with different prognosis and treatment strategies: luminal (ER+, PR+/-, HER2-); HER2+ (ER+/-, PR+/-, HER2+) and triple-negative breast cancer (TNBC; ER-, PR-, HER2-) [5]. ER+ and HER2+ patients are treated with targeted therapies against these biomarkers, including hormonal therapies like tamoxifen or HER2 inhibitors. On the contrary, TNBC patients are usually prescribed chemotherapies since targeted therapies are still unavailable [6]. Since the adoption of this strategy, breast cancer patient outcomes have improved. However, there is still a high number of treatment failure cases and relapses, especially in TNBC patients [7].

Several cancer cell subpopulations coexist within a single lesion and display different epigenomics, genomics and transcriptomics profiles. This variability, usually referred to as intratumor heterogeneity (ITH), reflects the existence of distinct tumour subpopulations that may harbour different sensitivities to anticancer therapies [8] and have been linked to treatment failure and relapse [9]. Until recently, ITH research has mainly focused on cancer cells and their genomic and gene expression variability, thus studying ITH at the subclone or transcriptional cell state levels [10,11]. In the last few years, the role of the tumour microenvironment (TME) on ITH and response to treatment has been acknowledged. The TME, which aggregates cellular and noncellular components, such as immune cells or the extracellular matrix (ECM) [12], influences transcriptomics changes in the tumour compartment. At the same time, the tumour hijacks the TME to favour its own growth [13]. Thus, this active cross-talk within the tumour ecosystem is a source of selective pressure for cancer evolution and, ultimately, a key factor in treatment response [14]. Despite recent efforts, there is a lack of studies relating drug response with the spatial organisation of the tumour and the cross-talk with the TME.

Single-cell technologies, particularly single-cell RNA-seq (scRNA-seq), have become a prominent method for studying ITH and TME. scRNA-seq methods require sample disgregation, causing cell stress [15] and death and not preserving the spatial organisation of the tissue. However, carcinomas grow and evolve within a spatial context [16], so studying the microanatomical niches within a tissue can illuminate unexplored ITH sources. In this scenario, an incipient sequencing-based technology known as spatial transcriptomics (ST) is becoming more relevant since it maps high-resolution transcriptomics data on top of tissue slides, allowing the study of the distribution of cell types and their cell-to-cell communications, the border between the tumour and TME compartments and the localisation of different tumour subpopulations [17].

In this work, we acquired ST data from 9 patients with invasive adenocarcinomas stratified into luminal, HER2+ or TNBC subtypes. We predicted the sensitivity to >1,200 drugs to explore the therapeutic heterogeneity of breast cancer while accounting for the spatial context and the interaction between the tumour and TME compartments. For this purpose, we have exploited a new version of Beyondcell [18], a tool for identifying tumour cell subpopulations with different drug response patterns in scRNA-seq data that now allows users to work with 10x Visium ST data.

## Methods

### Visium ST data preprocessing

We created Seurat v4.3.0.1 [19] objects from raw counts, the corresponding metadata and tissue images. Spots marked as “Artefact” were filtered out, and we removed low-quality spots that did not meet any of the thresholds specified in Supplementary Table S1. Next, we normalised the filtered raw counts with the *SCTransform* function (*vst.flavor=“v1”*). We assessed the cell cycle effect using the *CellCycleScoring* function and plotting a Principal Component Analysis (PCA; *npcs=50*) coloured by *Phase* labels, as suggested in Seurat vignettes. Based on this plot, we regressed the cell cycle when needed using the *SCTransform* function with *vars.to.regress=c("S.Score", “G2M.Score”)*. In multi-region patients (CID44971, CID4465 and CID4535), the effect of the region was also regressed.

### Spot deconvolution

We used Robust Cell Type Decomposition (RCTD) v2.2.1 [20] to predict the proportion of different cell types contained within each spot. First, we constructed 3 subtype-specific deconvolution references using annotated scRNA-seq data coming from 26 primary tumours (11 luminal, 5 HER2+ and 10 TNBCs) [5]. Labelled cell types comprised B cells, cancer-associated fibroblasts (CAFs), cancer cells, endothelial cells, myeloid cells, normal epithelial cells, plasmablasts, perivascular-like (PVL) cells and T cells. As part of quality control preprocessing, we removed single cells that met any of the following criteria: i) cells with a percentage of transcripts mapping to mitochondrial genes >=15%; ii) cells with a percentage of transcripts mapping to ribosomal genes >=40%; iii) cells with a number of transcripts <=250 or >=50,000; iv) cells with a number of expressed genes <=200 or >=7,500. Then, we merged the scRNA-seq datasets by breast cancer subtype and input the filtered raw counts to the *Reference* function to compute subtype-specific deconvolution references using a maximum of 500 cells per cell type. Finally, we deconvoluted the spots in each tissue slide using the corresponding reference, the filtered raw counts, the coordinates and the default parameters of *create.RCTD* and *run.RCTD* functions.

### Estimation of tumour purity and spot categorization

To estimate the tumour purity of each spot, we computed ESTIMATE (Estimation of STromal and Immune cells in MAlignant Tumours using Expression data) v1.0.13 scores [21] using the SCTransform-normalised and regressed expression counts obtained with Seurat v4.3.0.1. We scaled this score, which is inversely proportional to tumour purity, between 0 and 1 across all spots in the sample.

Then, we categorised each spot as either tumour or TME based on the agreement of three annotation sources: i) the histopathological annotations when available (spots labelled as “Ductal carcinoma *in situ*” or containing the “cancer” keyword), ii) the scaled ESTIMATE score (<=0.4), which is inversely proportional to tumour purity, and iii) the proportion of deconvoluted cancer cells (>=0.6) in each spot. The spots that did not meet at least 2 of these criteria were labelled as TME and assigned the non-malignant cell type with maximum deconvoluted proportion.

### Subclone inference

We determined sample-wise clonal structures from inferred copy-number alteration (CNA) profiles using Single CEll Variational Aneuploidy aNalysis (SCEVAN) v1.0.1 [22]. Briefly, we input filtered raw counts to *pipelineCNA* function (*beta_vega=0.5, SUBCLONES=TRUE, ClonalCN=TRUE, plotTree=TRUE*) using the spots labelled as TME as reference. In this process, some TME spots were relabelled as tumour if their inferred CNA profile was similar to other tumour spots. Moreover, tumour spots with no inferred CNAs were marked as diploid.

### Breast Sensitivity Signature Collection generation

The Breast Sensitivity Signature Collection (SSc breast) contains 1,372 gene signatures that reflect the transcriptional differences between sensitive and resistant breast cancer cell lines before drug treatment. In order to obtain these signatures, we performed a differential expression analysis against the area under the curve (AUC) with limma v3.54.0 [23] for all compounds tested in at least 10 different breast cancer cell lines. We selected the top 250 up- and down-regulated genes in sensitive versus resistant cancer cell lines, ranked by the t-statistic, to create bidirectional gene signatures of 500 genes each. The AUC was used to measure drug response because, contrary to IC50, it can always be estimated without extrapolation from the dose-response curve and has shown more accuracy in predicting drug response [24].

Expression and drug response data were retrieved from three independent pharmacogenomics assays: the Cancer Therapeutics Response Portal (CTRP) v2 [25–27], the Genomics of Drug Sensitivity in Cancer (GDSC) v2 [28–30] and the PRISM [31,32] repurposing compendium through the DepMap portal v22Q4 [33]. As these sources are independent, several signatures refer to the same compound. Consequently, the 1,372 transcriptomic signatures that form the SSc breast reflect the predicted response to >1,200 drugs.

To verify that the cancer cell lines included in the signature generation analyses were, in fact, representative of breast cancer patients, we used the corrected lineage reported in the Celligner project [34], which provides a framework to align cancer cell lines to human tumours from large cohorts of human patients such as the Cancer Genome Atlas (TCGA).

### Beyondcell Score calculation and correlation

Beyondcell Scores (BCS) constitute a spot-wise enrichment metric that is normalised to penalise spots with many zeros and/or outliers (genes whose expression is much higher than the rest of the genes). A positive BCS indicates enrichment in the up-regulated genes that comprise the signature. Conversely, a negative BCS designates spots enriched in the down-regulated genes. BCS close to 0 indicate uncertainty on the enrichment directionality.

As the SSc breast collection was computed comparing treatment-naive sensitive versus insensitive cells, the BCS reflects the predicted sensitivity (normalised BCS > 0) or insensitivity (normalised BCS < 0) to the corresponding drug. Similarly, a normalised BCS > 0 indicates enrichment in the phenotype represented by a functional signature.

BCS were computed using the SCTransform-normalised counts and the *bcScore* function (*expr.thres=0.1*) from the beyondcell v2.2.0 package [18]. Not available (NA) values in the normalised BCS matrix were transformed to 0. Then, we combined all patient-wise results into a single object and regressed the effect of patient identity with *bcRegressOut* function.

To identify groups with similar drug sensitivity, we computed a vector with the mean BCS per drug signature for each patient and i) major TC (for all spots) or ii) tumour TC (for tumour spots), excluding groups with less than 50 spots or mixed TCs. Then, we calculated the Pearson correlation coefficients between vectors and clustered the results according to Ward’s method.

### Therapeutic cluster computation

We computed therapeutic clusters (TCs) with the *bcUMAP* function from the beyondcell v2.2.0 package. First, this function performs a dimensionality reduction via PCA (*npcs=50*). Then, it constructs a K-Nearest Neighbors (KNN) graph (*k.neighbors=20*) based on the Euclidean distance in PCA space. The top principal components (*pc*) for clustering are determined by drawing an elbow plot. Then, the cells are grouped using the Louvain algorithm, specifying different resolutions (*res*). The TCs and the UMAP projection computed by *bcUMAP* can be visualised with the *bcClusters* function.

We selected a *res=0.5* and *pc=20* for all spots and a *res=0.15* and *pc=10* for tumour spots.

### Neighbourhood analysis

For each patient, we selected the spots located at the inner and outer edge of the mixed TCs using the *RegionNeighbors* function (*mode="inner_outer"*) from the semla v1.1.6 package [35]. Then, we performed a neighbourhood enrichment analysis to assess whether the spots belonging to two major TCs co-localized more than expected by chance. *RunNeighborhoodEnrichmentTest* transforms the Visium data in a network, with each spot constituting a node with a maximum of six edges with other spots. For each pair of major TCs, this function randomly permutes the class labels a fixed number of times (*n.perm=1000*), calculates a null distribution with the number of edges between spots from different classes and returns a z-score computed with the number of observed edges and the mean and standard deviation of the null distribution. A z-score around 0 can be interpreted as random spot localization. A positive z-score indicates an over-representation of the label pair co-localization. In contrast, a negative z-score can be viewed as a spatial repellent effect of the label pair. Finally, we aggregated the results across samples by computing the mean z-scores of each major TC pair.

### Radial distances

We computed the radial distances to the centre of the tumour-rich TCs, which we designated as the tumour core, with the *RadialDistance* function from the semla v1.1.6 package. This metric is calculated from the border of the region of interest (ROI). Thus, radial distances are equal to 0 at the tumoural margin, and they become more positive or negative further from or closer to the tumour core, respectively.

We tested the correlation between functional or drug sensitivity BCS and the radial distance to the tumour core in each sequenced region. Negative Pearson correlation coefficients indicate increased functional enrichment or drug sensitivity at the proximity of the tumour core. In contrast, negative correlation coefficients reflect increased functional enrichment or drug sensitivity at the tumour periphery. The p-values were adjusted using False Discovery Rate (FDR) correction for multiple testing.

### Functional enrichment analysis

To compare spots from different patients, we first integrated all Seurat objects using SCTransform-normalised expression counts. Then, we performed a differential expression analysis with the log-normalised counts of the groups of interest (major TCs, tumour TCs or expression clusters from patient V19L29). We compared each group against the rest using the *FindMarkers* function (*min.pct=0, logfc.threshold=0*) from the Seurat v4.3.0.1 package, which implements a non-parametric Wilcoxon rank sum test. With the specified parameters, this function returns the log fold-change of the average expression of each gene between the two groups, which was used to rank all genes in the expression matrix. For each comparison, we performed a pre-ranked gene set enrichment analysis (GSEA) with the *fgsea* function (*minSize=15, maxSize=500, nPermSimple=10000*) from the fgsea v1.26.0 package [36] using the Hallmark and Reactome v2023.1 collections [37–39], the cancerSEA gene sets [40] and other functional signatures [41–47].

### Differential sensitivity analysis

To determine the drugs that specifically target the groups of interest (non-mixed major TCs or tumour TCs), we compared each group against the rest using the *bcRanks* function from the beyondcell v2.2.0 package. For each comparison, this function ranks the drugs in the SSc breast collection according to the residual’s mean and the switch point (SP). The SP is a signature-specific Beyondcell metric that reflects the directionality and homogeneity of the drug response throughout the experiment. An SP=0 indicates a homogeneously sensitive response to the drug, whereas an SP=1 indicates a homogeneously resistant phenotype. Intermediate SPs represent a heterogeneous response.

We kept the drugs with intermediate switch points (between 0.4 and 0.6) and lowest or highest residual’s mean (5th percentile and 95th percentile for major TCs; 1st percentile and 99th percentile for tumour TCs). These drugs display a heterogeneous sensitivity pattern across the experiment and are the most specifically effective or ineffective against the group of interest.

### Cell-cell communication analysis

We inferred cell-cell ligand-receptor interactions using CellChat v2.1.0 [48]. We created a spatial CellChat object with the SCTransform-normalised expression counts, metadata and coordinates from all patients, grouping the tumour spots by TCs and the TME spots by cell type. Then, we performed a cell-cell communication analysis using the Secreted Signaling, ECM-Receptor and Cell-Cell Contact databases with the *computeCommunProb* function (*type="truncatedMean", trim=0.1, distance.use=FALSE, interaction.range=250, contact.knn=TRUE, contact.knn.k=6, population.size=TRUE*). We filtered out results for groups with <10 spots and retrieved the significant (*thres=0.05*) interaction strengths for ligand-receptor pairs of interesting signalling pathways (*signaling=c("PD-L1", "IGF", "DESMOSOME")*) and cell groups (*sources.use=targets.use=c("TC1.1", "TC1.2", "TC2", "TC3", "T cells", "CAFs", "B cells")*) using the *subsetCommunication* function. Finally, we scaled the interaction strengths of each ligand-receptor pair between 0 and 1.

### Tumoural ROI definition

At resolution 0.25, we computed 9 gene expression clusters using SCTransform-normalised expression counts, *FindNeighbors(dims=1:30)* and *FindClusters* functions from Seurat v4.3.0.1 package. All expression clusters but number 7 were confined within unique spatial regions. Thus, we subdivided cluster 7 into two subclusters using the *FindSubCluster(resolution=0.075)* function. We defined 8 initial ROIs from the expression clusters overlapping globular tumour ducts (4, 5, 6, 7_1, 7_2, 8 and 9). Next, we refined each ROI by splitting spatially disconnected spots using the *DisconnectRegions* function from the semla v1.1.6 package and keeping the main group of connected spots as ROI. Finally, we selected ROIs with at least 100 spots from any subclone.

For each tumoural ROI, we defined the edge and inner regions with the *RegionNeighbors* function (*mode="inner"*) from the semla v1.1.6 package. The spots detected at the inner border of each ROI were labelled as "edge", whereas the rest were categorised as "inner". Then, we tested the differential TC proportion between the edge and inner regions for each ROI and subclone using Fisher’s exact test. The p-values were adjusted using FDR correction for multiple testing.

### Statistical analyses and visualisation

All statistical analyses were carried out using the R v4 programming language [49] and the rstatix v0.7.2 [50] package. Data manipulation and visualisation was performed using functions included in the tidyverse v2.0.0 [51], GenomicRanges v.1.50.0 [52], ggpubr v0.6.0 [53], ggseabubble v1.0.0 [54], ggsankey v0.0.99999 [55], eulerr v7.0.0 [56], SPOTlight v1.4.1 [57], ComplexHeatmap v2.16.0 [58,59], circlize v0.4.15 [60], corrplot v0.92 [61], RColorBrewer v1.1-3 [62], viridis v0.6.4 [63], patchwork v1.1.3 [64] and figpatch v0.2 [65] packages.

## Results

### Tumour and TME dissection in ST breast cancer samples

In order to spatially dissect the therapeutic and functional heterogeneity in breast cancer samples, we analysed 10x Visium ST data retrieved from public repositories [5,66–69] coming from 9 breast cancer patients stratified into luminal (n=2), HER2+ (n=3) and TNBC (n=4) subtypes. Of these 9 patients, 7 had histopathological annotations, and sample V19L29 (HER2+) contained two consecutive slides. In addition, 2 TNBC patients (CID4465 and CID44971) and 1 ER+ patient (CID4535) had 2 or 3 regions sequenced into the same slide. To classify each spot as either tumour or TME, we used four independent annotation sources with high overlap (Figure 1A−B): the pathologist annotations (Figure 1C), the ESTIMATE score, which is inversely proportional to the tumour purity (Figure 1D), the proportion of cancer cells per spot (Figure 1E) and the clonal composition of each sample based on copy-number alteration (CNA) profiles (Figure 1F). In total, we successfully annotated 18,207 malignant and 11,858 non-malignant spots (Figure 1G, Supplementary Figure S1A) coming from all 9 patients. Notably, deconvolution analysis revealed that epithelial cancer cells, cancer-associated fibroblasts (CAFs) and myeloid cells were present in all samples (Supplementary Figure S1B−D).

**Figure 1.**
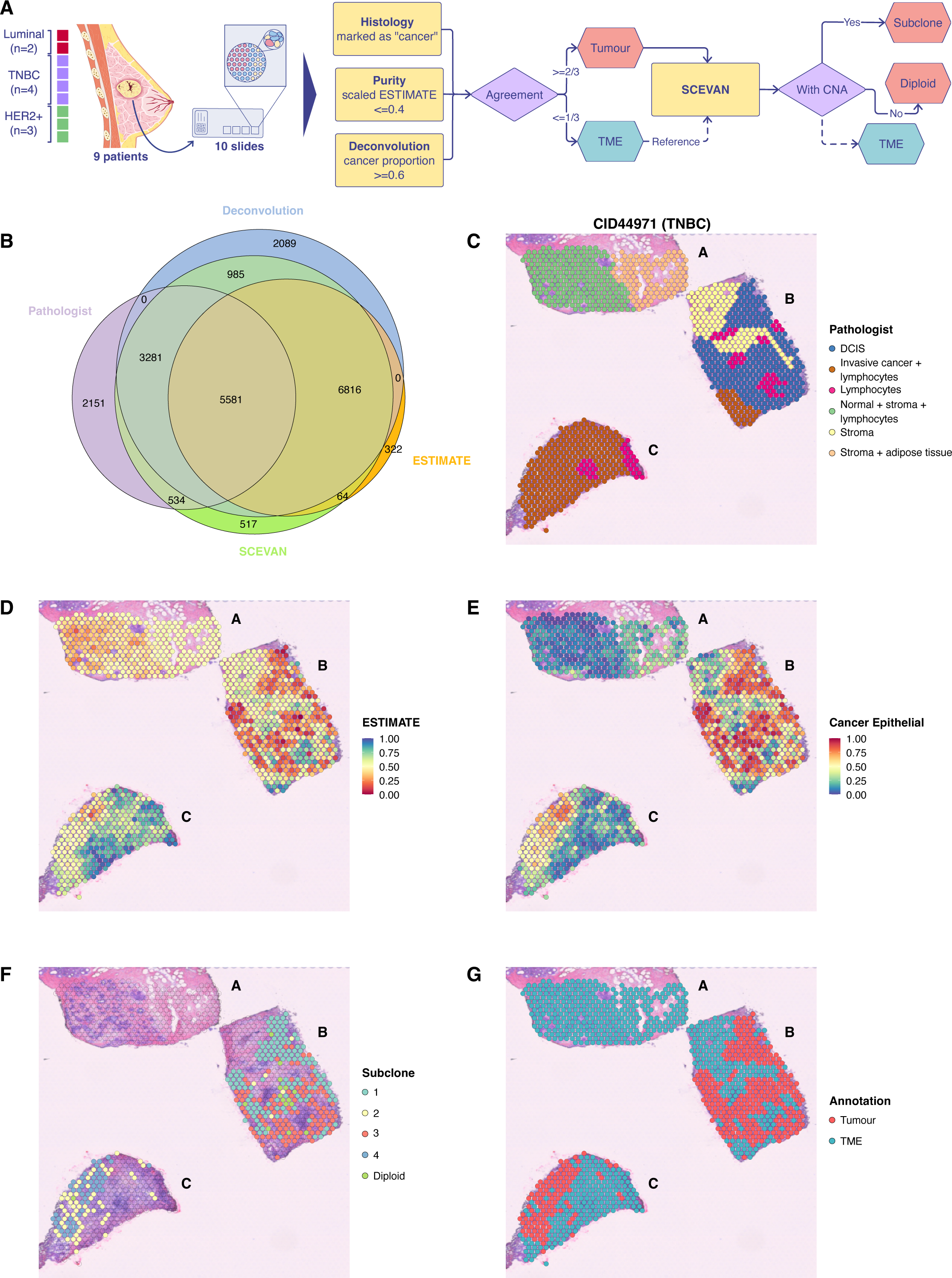
Tumour and TME dissection in ST breast cancer samples. **a)** Schematic representation of the analysis workflow followed to label each spot as either tumour or TME. **b)** Venn diagram representing the overlapping between the 4 annotation sources used to label tumour spots. An example tissue slide containing 3 regions coloured by **c)** pathologist annotations, **d)** scaled ESTIMATE score, which is inversely proportional to the tumour purity, **e)** cancer cell proportions and **e)** clonal structure of the tumour. **f)** Final tumour and TME labelling based on all previous annotations, as summarised in **a)**. **TME:** Tumour microenvironment; **ST:** Spatial transcriptomics; **ESTIMATE:** Estimation of STromal and Immune cells in MAlignant Tumours using Expression data; **TNBC:** Triple-negative breast cancer; **HER2+:** Human epidermal growth factor receptor 2 positive; **SCEVAN:** Single CEll Variational Aneuploidy aNalysis; **CNA:** Copy-number alteration; **DCIS:** Ductal carcinoma *in situ*.

### Spatial compartmentalisation of drug response in the tumour ecosystem

To predict the drug sensitivity of each spot, we created the Breast Sensitivity Signature Collection (SSc breast; Methods), which contains 1,372 gene signatures that reflect the transcriptional change between treatment-naive sensitive and resistant breast cancer cell lines to >1,200 compounds. With Beyondcell [18], we computed a matrix of enrichment scores for each spot and signature that was subsequently used to obtain 16 therapeutic clusters (TCs) with different predicted drug sensitivities (Figure 2A).

**Figure 2.**
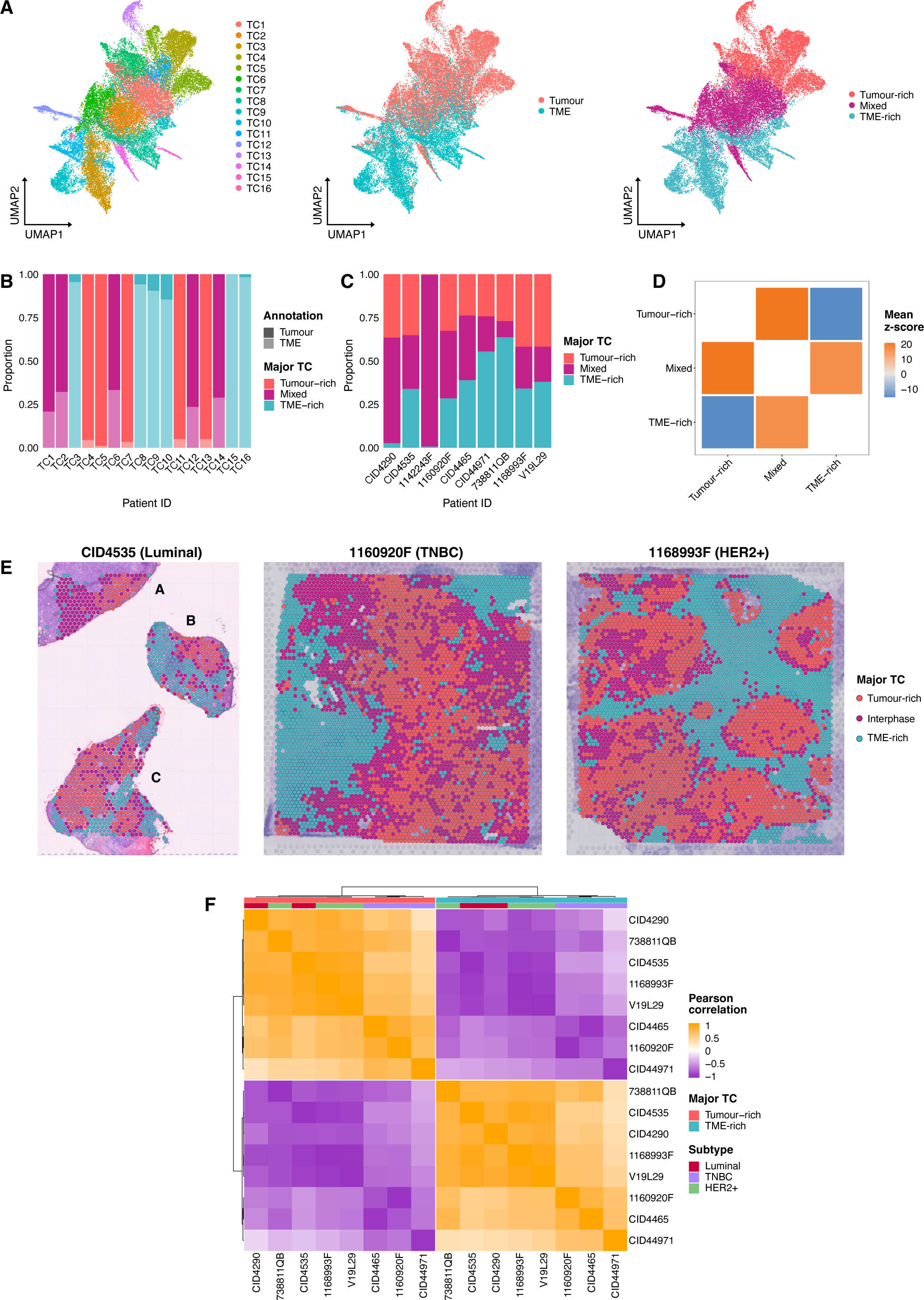
Spatial compartmentalisation of drug response in the tumour ecosystem. **a)** UMAP projection of spots from 9 breast cancer patients clustered according to their predicted sensitivity to the SSc breast collection. The same projection is coloured by TC (left), tumour or TME labels (centre) and major TC (right). **b)** Proportion of tumour and TME spots in each TC, coloured by major TC. The proportion of tumour spots was at least 95% in tumour-rich TCs, greater than 65%, less than 80% in mixed TCs, and 15% or less in TME-rich TCs. **c)** Proportion of the 3 major TCs across all patients. **d)** Neighbourhood enrichment between the spots defining each major TC’s edge. A z-score>0 indicates that the spots of two categories co-localize more than expected by chance, whereas a z-score<0 indicates a repellant effect between categories. Mean z-scores are computed to aggregate the information coming from all patients. **e)** Spatial projection of the 3 major TCs on top of tissue slides from different breast cancer subtypes. **f)** Correlation heatmap between the mean BCS of tumour and TME-rich compartments across all patients. Pearson correlation coefficients are clustered using Ward’s method. **UMAP:** Uniform Manifold Approximation and Projection; **SSc breast:** Breast Sensitivity Signature Collection; **TCs:** Therapeutic clusters; **TME:** Tumour microenvironment; **BCS:** Beyondcell Scores; **TNBC:** Triple-negative breast cancer; **HER2+:** Human epidermal growth factor receptor 2 positive.

We defined three major groups of TCs according to the observed proportion of tumour spots: tumour-rich (tumour proportion >=0.95), mixed (tumour proportion >0.65 and <0.80) and TME-rich (tumour proportion <=0.15) (Figure 2A−B). Interestingly, in mixed TCs, tumour and TME spots were grouped together, suggesting that therapeutic heterogeneity is not entirely driven by cell type identity (Supplementary Table S2). We confirmed that these three major TCs were present in all patients, implying similar drug responses independent of the breast cancer subtype (Figure 2C, Supplementary Table S3).

Next, we studied the spatial organisation of these major TCs. We observed that mixed TCs tended to co-localise with both tumour-rich and TME-rich TCs. Moreover, we noticed a clear repellent effect between spots in tumour-rich and TME-rich TCs (Figure 2D−E). These results indicate that the drug responsiveness defined by these major TCs is spatially organised and might overlap with the main compartments of the tumour ecosystem: the tumour and the TME regions and the interphase that physically separates them.

To assess whether drug response patterns were conserved across patients, we generated a correlation matrix from the Beyondcell Scores (BCS) within the tumour and TME regions of each patient (Figure 2F). Interestingly, these two spatial regions displayed opposite drug response patterns within the same patient, as indicated by their high anti-correlation, revealing the existence of intratumor therapeutic heterogeneity. At the same time, we identified two correlation clusters that perfectly matched our tumour and TME-rich annotations, suggesting recurrent response patterns within the same region across the whole cohort. Nevertheless, within the same cluster, the correlations that involved TNBC samples were lower than the rest, implying a higher interpatient therapeutic heterogeneity in TNBC samples.

Altogether, these results point to the existence of therapeutic heterogeneity between the spatial compartments of the tumour ecosystem. These different drug response patterns are conserved in all breast cancer patients independently of the subtype. Nevertheless, interpatient therapeutic heterogeneity exists and is particularly evident in TNBC tumours, in concordance with previous knowledge [70].

### A functional and drug sensitivity gradient exists between the tumour and TME compartments

In order to functionally characterise the major TCs, we first dissected their cellular composition and confirmed that the tumour-rich compartment was almost exclusively formed by cancer cells, together with a minority of CAFs and myeloid cells (Figure 3A). The interphase also contained endothelial cells, while the TME region displayed the highest cellular diversity. Overall, T cells appeared in the interphase and TME compartments, but interestingly, patient CID4465 (luminal) showed lymphocyte infiltration in the tumour region.

**Figure 3.**
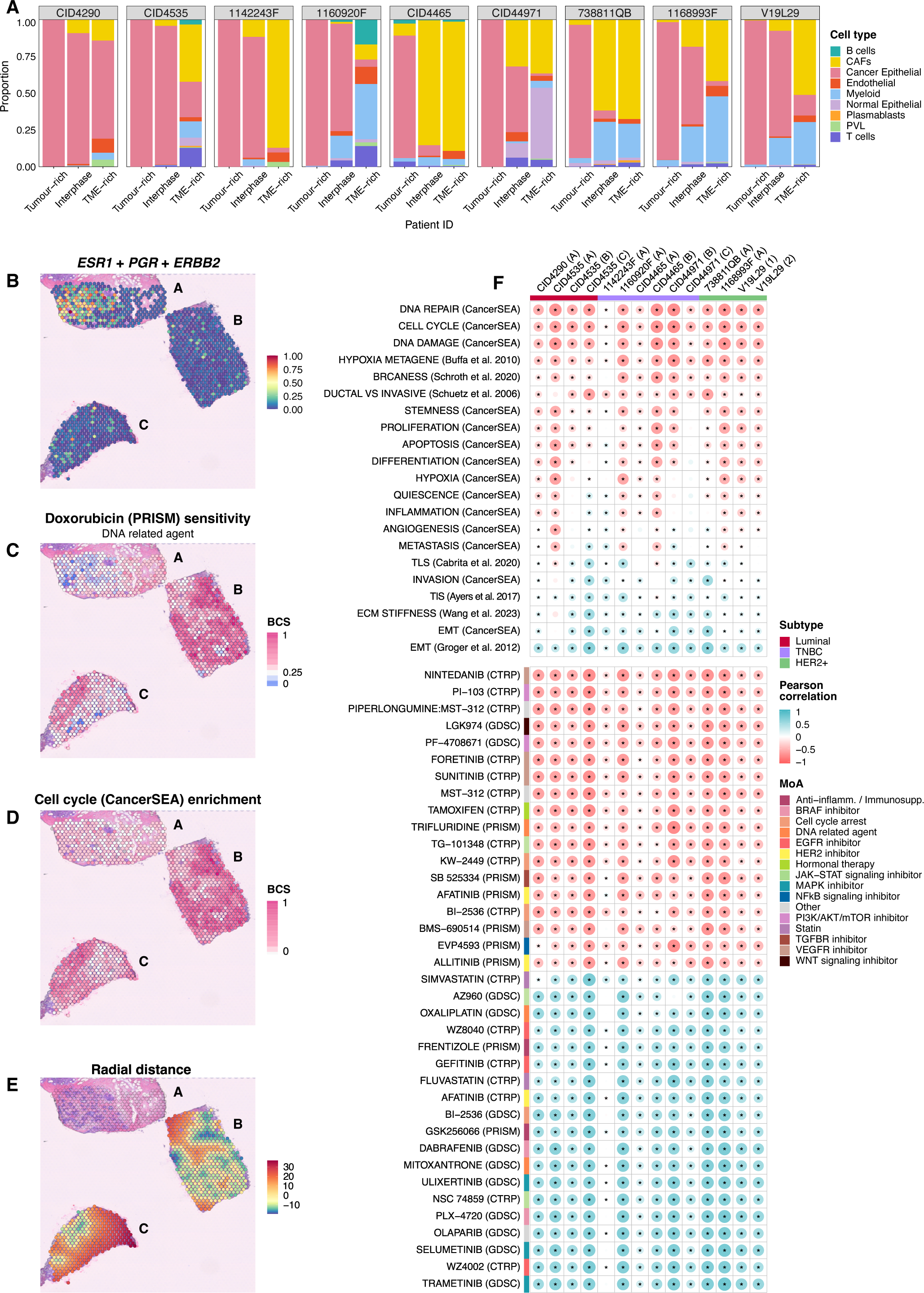
A functional and drug sensitivity gradient exists between the tumour and TME compartments. **a)** Proportion of cell types in each major TC across all patients. We labelled tumour spots as cancer cells and assigned TME spots to the non-malignant cell type with maximum deconvolved proportion. Spatial projection of **b)** breast cancer subtype biomarkers, **c)** drug sensitivity, **d)** functional BCS and **e)** the radial distances to the tumour core on top of patient CID44971 (TNBC) slide. To help visualisation, we plot the square root of radial distances multiplied by the original sign. **f)** Region-wise Pearson correlation coefficients between radial distances and functional (top) or sensitivity (bottom) BCS. Anti-correlated signatures are more enriched in spots closer to the tumour core. Correlated signatures are more enriched in the regions farthest away from the centre of the tumoural mass. The p-values were adjusted using FDR correction for multiple testing (*FDR<0.05). **TME:** Tumour microenvironment; **TCs:** Therapeutic clusters; **BCS:** Beyondcell Scores; **TNBC:** Triple-negative breast cancer; **FDR:** False Discovery Rate; **CAFs:** Cancer-associated fibroblasts; **PVL**: Perivascular-like cells; ***ESR1*:** Oestrogen Receptor gene; ***PGR*:** Progesterone Receptor gene; ***ERBB2*:** Erb-B2 Receptor Tyrosine Kinase 2 gene; **TLS:** Tertiary lymphoid structure; **TIS:** Tumour inflammation signature; **ECM:** Extracellular matrix; **EMT:** Epithelial to mesenchymal transition; **HER2+:** Human epidermal growth factor receptor 2 positive; **Anti-inflamm.:** Anti-inflammatory; **Immunosupp.:** Immunosuppressor.

We applied Beyondcell to compute enrichment scores for expression biomarkers (Figure 3B), cancer-related drugs like doxorubicin (Figure 3C) and well-known tumoral biological processes such as cell proliferation (Figure 3D). Taking advantage of the spatial information associated with each spot, we also calculated the radial distance to the core of the tumour-rich compartment (Figure 3E). Positive radial distances correspond to spots far away from the tumour core, whereas negative radial distances indicate proximity to the centre of the tumoural mass, and a distance of 0 denotes the tumoural margin. We then computed the region-wise correlation between radial distances and functional or sensitivity BCS (Supplementary Table S4−S6), discovering a functional and therapeutic gradient from the tumour core to the periphery of the tumoural regions (Figure 3F).

Notably, the spots within the tumour region were enriched in a ductal-like phenotype and predicted to be sensitive to standard hormonal therapies and HER2 inhibitors such as tamoxifen, afatinib and allitinib. We also observed an enrichment in proliferation and stemness functions closer to the tumour core. Accordingly, this region was more sensitive to cell cycle arrest agents (BI−2536 and KW−2449), telomerase inhibitors (MST-312), PI3K/Akt/mTOR inhibitors (piperlongumine, PI−103, PF−4708671) and WNT inhibitors (LGK974). Moreover, the tumour core was more hypoxic and paradoxically more sensitive to VEGFR inhibitors (BMS−690514, nintedanib, foretinib and sunitinib) since hypoxia triggers the expression of *VEGF*, a key regulator of angiogenesis. Interestingly, we observed a decreased sensitivity to DNA-related agents and olaparib within the tumour region, despite displaying a higher enrichment in DNA damage and BRCAness. This lower sensitivity can be due to an enrichment in DNA repair functions, which may help cancer cells repair DNA damage efficiently, contributing to treatment resistance. Indeed, upregulation of DNA repair mechanisms and hypoxia have been linked to chemoresistance in breast cancer [71,72]. On the other hand, the tumour periphery was more enriched in an invasive phenotype, inflammation, ECM stiffness and epithelial to mesenchymal transition (EMT). Consistently, these spots displayed increased sensitivity to JAK-STAT signalling inhibitors, statins and immunosuppressants or anti-inflammatory agents, drugs that mainly target the immune component and the inflammatory response of the TME.

These functional and therapeutic differences were confirmed by compartment-wise functional analysis and drug ranking (Supplementary Figure S2A−B, Supplementary Table S6−8). We assessed that the interphase displayed medium sensitivities to all these drugs (mean BCS for Tumour-specific therapies: Tumour-rich=7.74, TME-rich=−7.27, Interphase=−0.52; mean BCS for TME-specific therapies: Tumour-rich=−7.92, TME-rich=7.44, Interphase=0.53), suggesting the existence of a therapeutic continuum from the tumour core to the TME and *vice versa*.

Using Beyondcell, we checked these predicted trends in three examples of breast cancer clinical subtypes. As expected, only a normal epithelial tissue patch in region A of the TNBC sample expressed *ESR1* (ER gene), *PGR* (PR gene) and *ERBB2* (HER2 gene) biomarkers (Figure 3B). The triple-negative tumour-rich region displayed specific sensitivity to first-line treatment doxorubicin (Figure 3C), which can be explained by an enrichment in cell cycle activity in the same region (Figure 3D). Conversely, in luminal and HER2+ samples, the high expression of *ESR1* and *ERBB2* biomarkers in the tumour-rich region was correlated with the predicted response to standard-of-care therapies like tamoxifen or HER2 inhibitors (Supplementary Figure S3A−C). Furthermore, for each TME spot, we computed a single sample bidirectional enrichment score [73] depicting its protumor (>0) or antitumor (<0) status, according to the BCS ranking of 22 pan-cancer microenvironment signatures [74]. We observed TME heterogeneity among samples (Supplementary Figure S3D). The TME of the luminal example was predominantly antitumoral and thus did not need chemical inhibition. In contrast, the HER2+ patient displayed a dual TME whose protumoral region could also be targeted with the HER2 inhibitor afatinib.

### The interaction with the TME influences cancer drug response

To determine whether there was therapeutic heterogeneity within the tumour region, we reanalysed the spots labelled as tumour with Beyondcell. We obtained 5 TCs (Figure 4A) and performed a correlation analysis (Figure 4B, Methods), identifying response pattern similarities between TC3 and the patient-specific clusters TC4 and TC5. Thus, we regrouped all of them into a new TC3. On the other hand, we noticed that TC1 could be subdivided into two subclusters with different response patterns. After this refinement process, we obtained 3 TCs, one divided into two subclusters (TC1.1 and TC1.2) (Figure 4C), that were conserved across patients (Supplementary Figure S4A, Supplementary Table S9).

**Figure 4.**
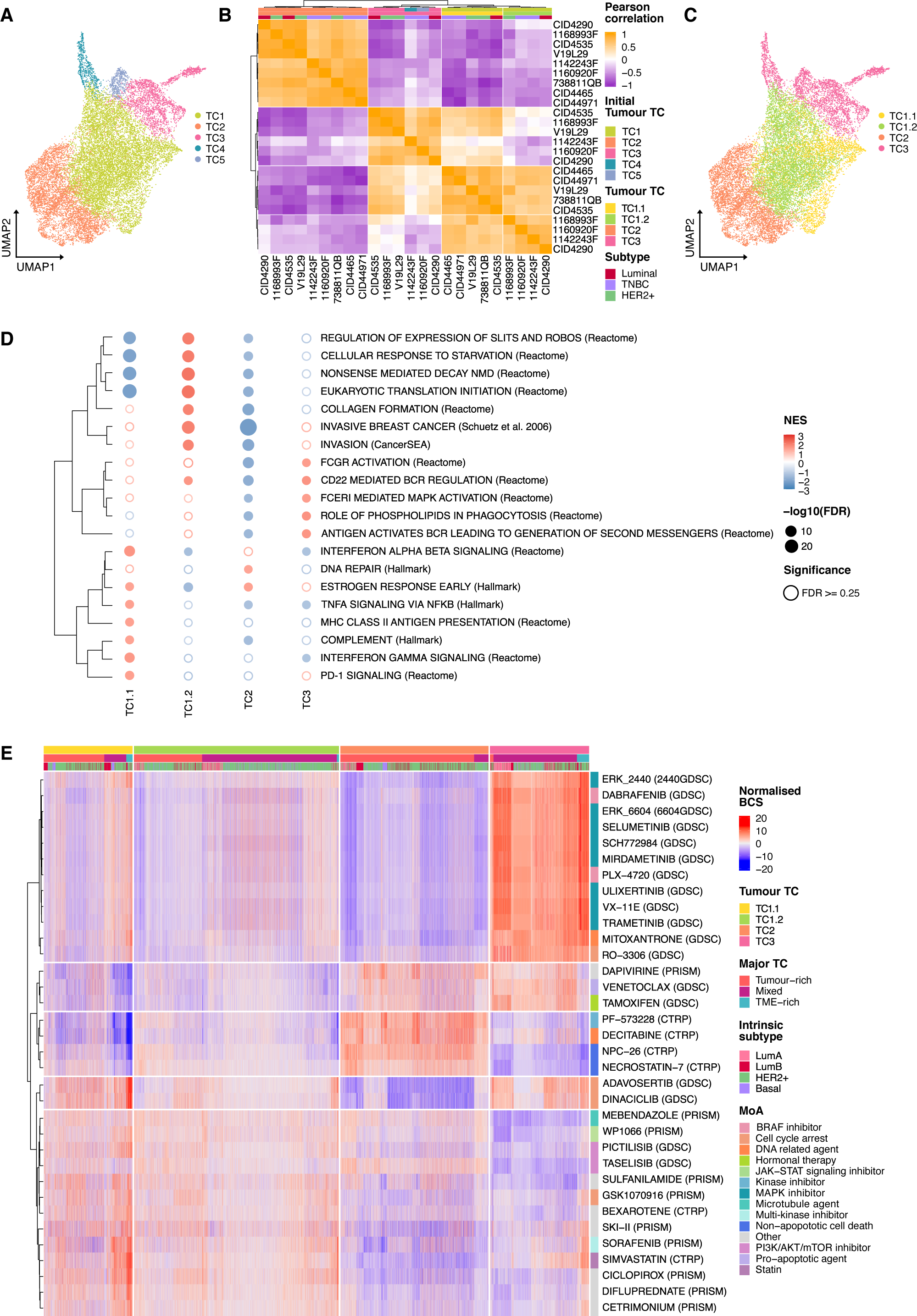
The interaction with the TME influences cancer drug response. **a)** UMAP projection of tumour spots from 9 breast cancer patients clustered according to their predicted sensitivity to the SSc breast collection, coloured by initial TC. **b)** Correlation heatmap between the mean BCS of tumour TCs across all patients. Pearson correlation coefficients are clustered using Ward’s method. According to this clustering, initial tumour TCs were refined into 3 TCs, one divided into two subclusters (TC1.1 and TC1.2). **c)** The same projection as in **a)**, coloured by refined tumour TCs. **d)** Bubble heatmap depicting significantly positively enriched pathways in each tumour TC, as identified by differential gene expression analysis and pre-ranked GSEA. Rows represent functional pathways, and columns represent the tumour TCs. The colour of the bubble is proportional to the NES magnitude and the size of the bubble to the FDR-adjusted p-value. Empty bubbles represent non-significant results (FDR>=0.25). Rows are clustered according to the Euclidean distance between NES. **e)** Heatmap of drugs that specifically target the tumour TCs. Drugs are clustered according to their BCS using Ward’s method and labelled by MoA. Spots from all samples are ordered by tumour TC, major TC and the intrinsic subtype determined using single-cell PAM50 signatures derived from Wu *et al.* **TME:** Tumour microenvironment; **UMAP:** Uniform Manifold Approximation and Projection; **SSc breast:** Breast Sensitivity Signature Collection; **TCs:** Therapeutic clusters; **BCS:** Beyondcell Scores; **GSEA:** Gene Set Enrichment Analysis; **NES:** Normalised Enrichment Score; **FDR:** False Discovery Rate. **MoA:** Mechanism of action; **TNBC:** Triple-negative breast cancer; **HER2+:** Human epidermal growth factor receptor 2 positive; **LumA:** Luminal A; **LumB:** Luminal B.

We verified that these TCs were different in terms of proximity to the tumour core and compartment affiliation (Supplementary Figure S4B−C). TC2 was the closest to the centre of the tumoural mass and displayed the highest percentage of tumour-rich spots (TC1.1=68%, TC1.2=33%, TC2=90% and TC3=4%). In contrast, TC3 was the more distal cluster, mainly composed of interphase spots (TC1.1=25%, TC1.2=66%, TC2=10% and TC3=84%). Thus, we hypothesised that TC2 constituted the tumour core, whereas TC3 comprised the tumoural margin.

Functional enrichment analysis (Figure 4D, Supplementary Table S6,S10) revealed that TC2 was the least invasive TC (e.g. downregulation of invasion gene set or collagen formation). Furthermore, ligand-receptor analysis (Supplementary Figure S4D) confirmed that TC2 spots were in close contact via DSC2-DSG2 interactions. These cadherins are primary constituents of the desmosome, a cell-cell junction that maintains the integrity of normal epithelia. Thus, these results convey that the tumour core represented by TC2 is in a pre-invasive state. The remaining TCs were enriched in functions that suggested communication with the TME. TC3 was characterised by B cell activation, phagocytosis and triggering of MAPK signalling. TC1.1 was defined by the triggering of the complement cascade, cytokine signalling (TNFα and IFNα, β and γ) and functions that indicated a T cell suppressive state via PD-1 signalling. Conversely, TC1.2 displayed an increased translational activity triggered in response to starvation, as well as upregulation of NMD, Slit/Robo signalling and collagen formation, which might be related to ECM reorganisation, contributing to its invasive phenotype. The upregulation of these functions points to nutrient deprivation due to inadequate blood supply, which may favour a hypoxic niche. Some authors have proposed intron retention, an aberrant splicing event which can introduce premature stop codons and lead to NMD, as a possible mechanism by which hypoxia can promote angiogenesis, cancer cell migration and invasiveness in breast cancer (Han et al. 2017). Ligand-receptor analysis (Supplementary Figure S4D) confirmed that TC1.1 interacted with suppressed T cells via PD1-PDL1 binding and TC1.2 with CAFs via IGF1R signalling, which has been related to invasiveness, EMT and resistance to EGFR inhibitors such as gefitinib and chemotherapies like gemcitabine and paclitaxel in breast cancer [75]. Collectively, these results imply that each TC exhibits distinct functional states conditioned by interactions with different components of the TME.

Using the SCSubtype gene signatures from Wu *et al.* [5], we classified the tumour spots into intrinsic subtypes (Luminal A, Luminal B, HER2+, or Basal) based on the maximum BCS, with ties resulting in an "Undetermined" annotation. Next, we identified drugs specifically targeting each tumour TC, but no TC was enriched in a particular intrinsic subtype (Figure 4E, Supplementary Table S11). The tumour core depicted by TC2 displayed sensitivity to non-apoptotic cell death agents and kinase inhibitors, whereas TC3 appeared to be differentially sensitive to MAPK inhibitors. Interestingly, both TC1 subclusters displayed lower sensitivity to their specific drugs compared with the other clusters (mean BCS for specific therapies: TC1.1=2.35, TC1.2=1.08, TC2=6.15, TC3=7.95). Still, cancer cells in an immunosuppressive microenvironment defined by PD1-PDL1 binding or interacting with CAFs via IGF1R signalling exhibited specific sensitivity to cell cycle arrest agents and PI3K/AKT/mTOR inhibitors. Our analysis also indicated that TC1.1 and TC1.2 were sensitive to SKI-II, a sphingosine kinase (SK) inhibitor and simvastatin. Up-regulation of sphingosine kinase 1 (*SK1*) has been associated with invasiveness and chemoresistance in breast cancers, and targeting SK1 could have therapeutic potential [76]. Moreover, simvastatin has been proposed to attenuate breast cancer metastases and recurrence [77,78]. Of note, none of these subclusters was predicted to be sensitive to the standard-of-care tamoxifen, whereas the rest of the TCs displayed high sensitivity to this drug. This result highlights the relevance of studying ITH to identify pre-resistant tumour subpopulations that, if overlooked, can lead to treatment failure and relapse. We hypothesised that the interacting TME influenced the low drug sensitivity observed in TC1. We confirmed that TC1.1 was significantly enriched in the tumour inflammation signature (TIS) gene set [44], which reflects a suppressed adaptive immune response and has been associated with sensitivity to anti-PD1 therapies [44,79] (Supplementary Figure S4E). On the other hand, TC1.2 was significantly enriched in ECM stiffness compared to the rest of the TCs (Supplementary Figure S4F). ECM stiffness is linked to CAF activity, constitutes a mechanical barrier for treatment diffusion and promotes treatment resistance by triggering different signalling pathways in tumour cells [80].

Altogether, these results support the idea that intratumoral therapeutic heterogeneity does exist in breast cancer, and some niches may display decreased sensitivity, or even insensitivity, to standard-of-care treatments. These differences may be related to distinct transcriptomics states triggered by the interaction with specific components of the TME, which can sensitise to immunotherapies or protect cancer cells from drugs.

### Subclones display heterogeneous responses depending on their location in tumour ROIs

We asked ourselves whether different subclones drove therapeutic heterogeneity within the tumour region. In order to explore this hypothesis, we selected patient V19L29 (HER2+) because it was the sample with more sequenced tumour spots (n=4,713; Supplementary figure S1A), granting higher statistical power. We defined 6 tumoural regions of interest (ROIs) based on well-delimited globular ducts (Figure 5A, Supplementary Figure S5A). We confirmed that ROIs had a biological identity, as they were identified as different expression clusters (Supplementary Figure S5B) enriched in different functions (FDR<0.25) such as EMT activation in ROI5 or hypoxia in ROI4 among others (Supplementary Figure S5C, Supplementary Table S6,S12). These ROIs contained 6 cancer subclones defined by CNA profiles (Figure 5B), grouped into TC1.1, TC2 and TC3 (Figure 5C). Subclones showed ROI-specificity (i.e. were localised in one or few ROIs), but the same subclone belonged to different TCs, suggesting intra-subclonal therapeutic heterogeneity (Figure 5D). We asserted that, for some subclones and ROIs, the proportion of TCs was significantly different (FDR<0.05) between the edge and inner regions (Figure 5E), TC2 being more abundant in the inner region and TC1.1 and TC3 overrepresented at the edge. This result implies that the spatial location of spots, even with the same genetic background, conditions their transcriptomic profile and might ultimately result in different response patterns, highlighting the importance of studying ITH in its spatial context.

**Figure 5.**
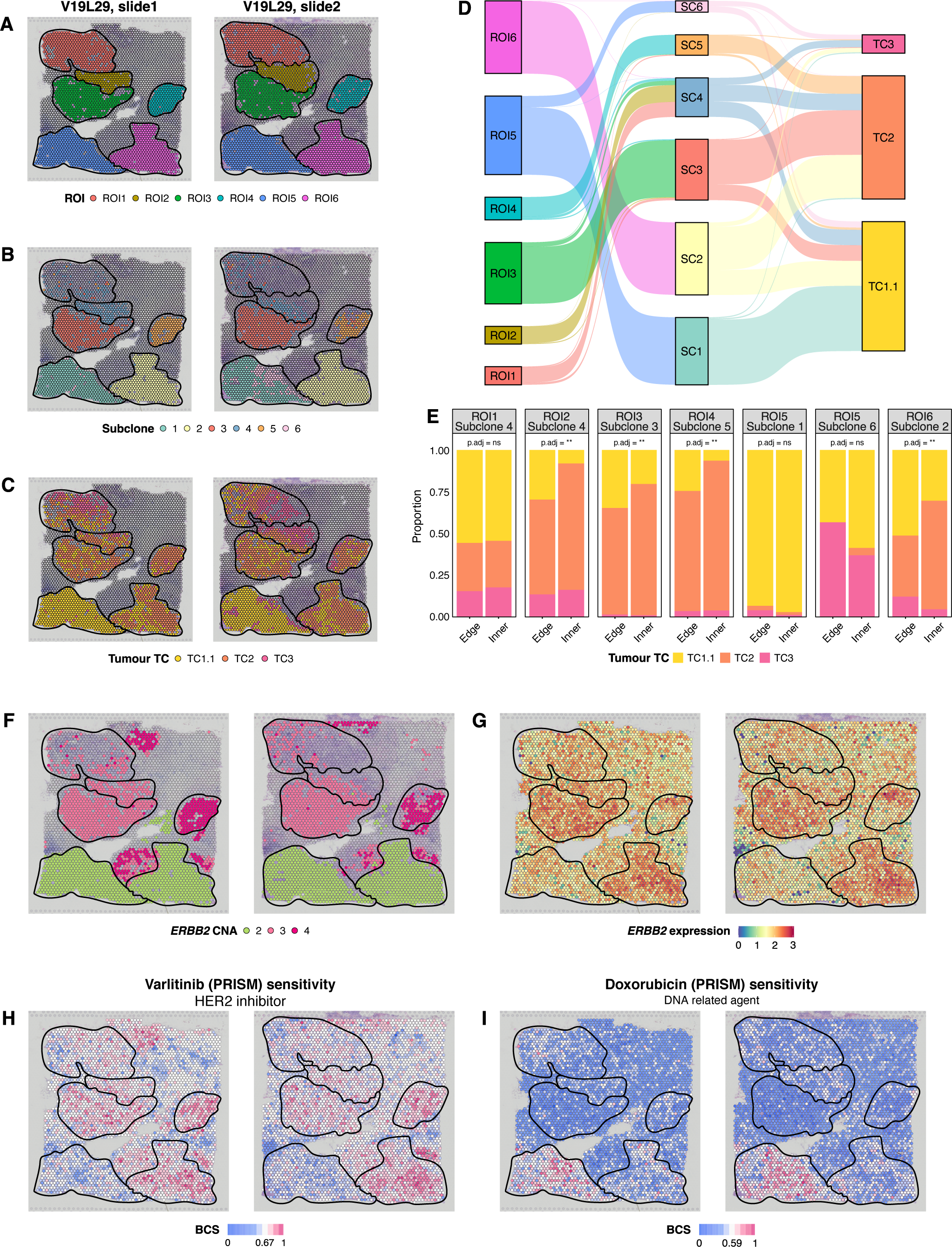
Subclones display heterogeneous responses depending on their location in tumoural ROIs. Spatial projection of **a)** ROIs, **b)** tumour subclones and **c)** tumour TCs on top of patient V19L29 (HER2+) tissue slides. **d)** Sankey diagram outlining spots’ ROI, subclone and TC membership. **e)** Barplots depicting the different TC compositions between the edge and inner regions of each subclone and ROI. Differences in proportions were tested with a Fisher’s exact test. The p-values were adjusted using FDR correction for multiple testing (ns FDR>=0.05, *FDR<0.05, **FDR<0.01, ***FDR<0.001, ****FDR<0.0001). Spatial projection of **f)** the CNAs and **g)** the expression for *ERBB2* biomarker, **h)** the sensitivity to the HER2 inhibitor varlitinib and **i)** the sensitivity to doxorubicin. This patient constitutes an example of pre-existent treatment resistance and the usefulness of dissecting ITH for therapy design. A combination of varlitinib and doxorubicin would target both HER2+ and non-HER2+ cancer cells to eliminate the tumour and avoid relapse. **ROI:** Region of interest; **TCs:** Therapeutic clusters; **HER2+:** Human epidermal growth factor receptor 2 positive; **FDR:** False Discovery Rate; **ns:** Not significant; **CNAs:** Copy-number alterations; ***ERBB2*:** Erb-B2 Receptor Tyrosine Kinase 2 gene; **ITH:** Intratumor heterogeneity; **SC:** Subclone; **BCS:** Beyondcell Scores.

In the clinic, the average overexpression or amplification of *ERBB2* guides the administration of HER2 inhibitors to HER2+ breast cancer patients. However, some authors have described molecular subtype heterogeneity between cancer cells within the same lesion [5]. Interestingly, subclones 1 and 6 contained in ROI5 were diploid for *ERBB2* and had a lower *ERBB2* expression than the rest of the tissue (Figure 5F−G). Accordingly, Beyondcell reported insensitivity to the HER2 inhibitor varlitinib in ROI5, whereas the rest of the tumour was predicted to be responsive (Figure 5H). If patient V19L29 were given HER2 inhibitors as monotherapy, ROI5 would survive to regenerate a HER2 inhibitor-resistant tumour. Beyondcell identified doxorubicin as a complementary therapy to target ROI5 specifically (Figure 5I). Thus, we propose a combination of varlitinib and doxorubicin to target the major subpopulation of HER2+ cancer cells and the minor non-HER2+ subclones to eliminate the tumour and avoid relapse. This case constitutes an example of pre-existent treatment resistance and the usefulness of dissecting ITH for therapy design.

## Discussion

One of the main challenges in personalised precision oncology is to tackle ITH to boost treatment effectiveness and avoid relapse [8,81]. Thus, this heterogeneity has been extensively studied to characterise cancer subpopulations across samples and identify specific drug vulnerabilities [18,82–84]. However, the impact of the TME on the tumour ecosystem has been usually overlooked. Current research indicates that the interactions with the TME are critical for tumour progression, favouring vascularisation, ECM remodelling, immune exclusion and suppression [85]. The selective pressure exerted by the local TME leads to the diversification of malignant and non-malignant cell subpopulations within the same tumour, thus enhancing ITH and increasing the chances of therapeutic resistance [86]. Therefore, increasing interest is in studying the tumour ecosystem and identifying therapeutic niches. Targeting specific elements of the TME holds the potential to reduce the incidence of metastasis, which is a major contributor to breast cancer mortality rates. Moreover, the TME components exhibit some conservation across breast cancer subtypes, providing a comprehensive therapeutic target [87]. Sequencing-based ST opens a new world of possibilities to study the organisation of tissues and the interactions between the cancer cells and the TME [88]. In this work, we have leveraged sequencing-based ST and Beyondcell [18] to identify therapeutic niches based on the sensitivity predictions to a collection of over 1,200 drugs. Our findings revealed a compartmentalisation of the drug response in breast cancer, unveiling a sensitivity and therapeutic gradient relative to the distance to the tumour core. Moreover, we dissected the therapeutic heterogeneity within the tumour region, which appears to be related to different TME interactions conserved across patients. Furthermore, we observed different response patterns within the same cancer subclone depending on its spatial location.

We reported enrichment in cell proliferation, stemness and hypoxia near the tumour core, whereas inflammation, invasiveness, and EMT were enriched in the periphery. This functional gradient could be a consequence of the co-existence of diverse molecular gradients, including oxygen, nutrients, pH, chemokines, and growth factors within the tumour ecosystem [89–92]. Most of these gradients result from the leaky vasculature of solid tumours and the rapid proliferation of cancer cells. Oxygen and nutrient concentration decrease with the distance to blood vessels. Thus, distant cells adapt to the limited oxygen and nutrient supply by triggering transcriptomics and metabolic changes that decrease their proliferation rate. In response to hypoxia, cells undergo anaerobic glycolysis, where pyruvate is converted to lactate and released to the extracellular space. Consequently, the medium becomes more acidic, creating a pH gradient. The secretion of growth factors and chemokines by TME cells generates additional biochemical gradients that induce functional changes in surrounding cells. Low oxygen concentrations, an acidic medium and secreted chemokines favour cell migration and EMT [93]. These molecular gradients and the transcriptional states they induce also influence drug susceptibility. Hypoxia has been associated with increased DNA repair activity and subsequently reduced efficacy of multiple chemotherapeutic agents. This phenomenon induces the expression of Multidrug Resistance Protein 1 (MDR1) and Breast Cancer Resistance Protein (BCRP), two ABC transporters that decrease the intracellular concentration of chemotherapeutic agents favouring resistance [94]. In addition, an acidic microenvironment can ionise weakly basic chemotherapies such as mitoxantrone, reducing their cellular uptake and cytotoxicity [95,96]. Accordingly, we observed an enrichment in DNA repair activity in the hypoxic tumour core and a decreased sensitivity to different chemotherapies, including mitoxantrone. Moreover, this hypoxic state could also explain the increased sensitivity to VEGFR inhibitors near the centre of the tumour. On the other hand, the enrichment in inflammation at the periphery of the tumour is coherent with an increased sensitivity to immunosuppressants and anti-inflammatory agents in this region. These sensitivity and functional gradients defined three therapeutic regions that overlapped with the three main compartments of the tumour ecosystem: the tumour, the TME, and the interphase that physically separated the previous two.

We also dissected the therapeutic heterogeneity of spots labelled as tumour and characterised the resultant TCs, which were conserved across patients, linking function and ligand-receptor interactions to drug response within the tumour compartment. The tumour core, characterised by a non-invasive phenotype and active metabolism, was predicted to be sensitive to kinase inhibitors and non-apoptotic cell death agents. In contrast, cancer cells close to B lymphocytes were specifically sensitive to MAPK inhibitors. The remaining TC was subdivided into subclusters with different biological functions and TME interactions. Cancer cells in TC1.1 inhabited an immunosuppressive microenvironment promoted by PD1-PDL1 interactions. The binding of the cancer ligand PD-L1 to its immune receptor PD-1 causes the suppression of cytotoxic response in activated CD8+ T cells, thus guaranteeing the survival of cancer cells. Consequently, the inhibition of this interaction by immune checkpoint inhibitors (ICI) shows great efficacy in tumours with this immune-suppressive phenotype [85]. We confirmed ICI sensitisation by the significant enrichment of TIS in TC1.1. On the other hand, TC1.2 displayed an increased collagen production and an invasive phenotype. This phenotype has been related to the presence of CAFs, which are known to cause matrix remodelling and promote invasiveness. Indeed, we confirmed that TC1.2 was interacting with CAFs via IGFR1 signalling. ECM remodelling can reduce the sensitivity to targeted therapies by i) creating a protective niche that physically impedes the delivery of drugs; ii) mechano-transduction of proliferation or anti-apoptotic signals mediated by integrins or iii) metabolic changes induced in cancer cells due to the internalisation of ECM components. These mechanisms have been observed in breast cancer *in vivo* and *in vitro* models, which displayed increased resistance to HER2 and multi-kinase inhibitors [97]. Accordingly, TC1.2 was significantly enriched in ECM stiffness, which may explain the overall low sensitivity of this subcluster to the entire collection of tested drugs. These results pinpoint how the interaction with different components of the TME can modulate the function of cancer cells and ultimately lead to greater therapeutic heterogeneity within the same patient.

Finally, based on the results of a single patient, we propose that the TME also modulates the expression of cancer cells with identical genetic backgrounds. Although we would need a higher sample size to extract robust conclusions, therapeutic heterogeneity in single-cell-derived clones has been previously described and associated with different cell states before drug administration [98]. Moreover, the same authors have proposed that this transcriptional heterogeneity may be rooted in responses to external cues. Thus, studying the subclonal composition alone without considering the interacting TME may not be enough to dissect and understand the therapeutic heterogeneity of tumours. Adaptive therapy, which seeks to make cancer a chronic disease by letting treatment-sensitive cells survive, compete and eliminate resistant subpopulations [99], or clonetherapy, which aims to use drug combinations against both the major cancer subpopulation and minor pre-resistant cells [100] could benefit from studies like our own. We expect that dissecting ITH and identifying recurrent programs involved in therapeutic response will guide more precise treatments and aid these novel therapeutic strategies to tackle drug resistance.

The findings of this analysis point out some potential improvements and considerations for the future. Spots in 10x Visium technology do not correspond to single cells but groups of 1 to 10 cells. Thus, researchers use a scRNA-seq reference to deconvolve the cell types contained in each spot and assign a unique label according to arbitrary proportion thresholds. Moreover, these spots are discrete, thus leaving a large portion of the tissue unmeasured. In this study, we categorised spots as tumour or TME based on the agreement of four highly overlapping annotation sources. Although the spots labelled as tumour contained a minor proportion of non-cancer cell types, we could detect ITH, therapeutic heterogeneity and even different subclones using tools developed for single-cell technologies. Nevertheless, to precisely apply these methods, we could leverage new technologies that allow for the sequencing of single cells in the spatial context, such as slide-tags [101] or Visium HD [102], or machine learning methods like TESLA [103], which generates pixel-level resolution gene expression profiles. Moreover, the spatial organisation of drug response that we observed encourages the use of spatial-informed clustering methods implemented in tools like Giotto [104], stLearn [105] or BayesSpace [106] in future analysis. In the present study, only one patient was sequenced in two consecutive adjacent sections, with virtually no heterogeneity between them. As the Visium capture area is only 6.5×6.5mm, several adjacent (horizontal) and serial (vertical) tissue sections would be necessary to create a three-dimensional reconstruction of the tumour and study the therapeutic differences between cancer cells in the core and the periphery of the tumour. Pioneering work in other biomedical areas [107] could serve as a reference in cancer research, which would benefit from tools developed for horizontal and vertical data integration like GraphST [108]. Moreover, we expect that novel ST-specific batch correction methods like PRECAST [109] would be handy for extracting insights from integrated data by performing spatial-informed clustering across samples. Still, observing spatial organisation from just expression clustering suggests that expression alone can explain most spatial variability. Thus, a potential application of these ST analyses is to extract geographic expression signatures that can be remapped onto scRNA-seq data to clarify expression differences between clusters of the same cell type. Furthermore, ITH is a multi-level phenomenon that includes genomics, epigenomics, transcriptomics, proteomics and the interaction with the TME. Multi-omics analysis, facilitated by the recent release of Seurat v5 [110], would further our understanding of ITH and its relationship with therapeutic response. Finally, our collection of drug signatures derives from screenings in cancer cell lines with no interactions with the TME. Moreover, we lack specific drugs to target the immune system or the stromal component. In this sense, our SSc breast collection would greatly benefit from signatures derived from drug screenings in co-cultures to determine the contribution of the TME to the drug action. On top of that, we must not overlook that non-cellular constraints such as matrix stiffness, low oxygen concentrations or pH variations influence treatment diffusion and efficacy. Thus, future efforts to characterise the effect of non-cellular TME components on cellular function and drug response would benefit therapeutic heterogeneity dissection in breast and other cancer types.

## Conclusions

In this work, we have leveraged Beyondcell to group sequencing-based ST data from 9 breast cancer patients according to the predicted sensitivity of each spot to a collection of >1,200 drugs. Our results highlight how drug response patterns are organised in space and match the different compartments of the tumour ecosystem. Moreover, we observed a therapeutic and functional gradient that evolves from the tumour core and identified tumour- and TME-specific drug sensitivities and biological pathways. Within the tumour compartment, we defined conserved niches with different response patterns depending on their interactions with the TME. We propose that the proximity to the TME may modulate drug response within the same cancer subclone. Thus, ITH and, ultimately, therapeutic heterogeneity would depend not only on the genetic background of cells but also on their transcriptional state, influenced by the interaction with the TME. We also provide use case examples of how Beyondcell can be used on spatial transcriptomics data to guide therapy selection for individual patients.

## List of abbreviations

AUC: Area under the curve
CAF: Cancer-associated fibroblast
CNA: Copy-number alteration
CTRP: Cancer Therapeutics Response Portal
BCS: Beyondcell Scores
DCIS: Ductal carcinoma *in situ*
ECM: Extracellular matrix
EMT: Epithelial to mesenchymal transition
ER: Oestrogen receptor
ESTIMATE: Estimation of STromal and Immune cells in MAlignant Tumours using Expression data
FDR: False Discovery Rate
GDSC: Genomics of Drug Sensitivity in Cancer
GSEA: Gene Set Enrichment Analysis
HER2: Human epidermal growth factor receptor 2
H&E: Hematoxylin-Eosin
ICI: Immune checkpoint inhibitors
ITH: Intratumor heterogeneity
KNN: K-Nearest Neighbors
MoA: Mechanism of action
NA: Not available
NES: Normalised Enrichment Score
PCA: Principal Component Analysis
PR: Progesterone receptor
PVL: Perivascular-like
RCTD: Robust Cell Type Decomposition
ROI: Region of interest
SCEVAN: Single CEll Variational Aneuploidy aNalysis
scRNA-seq: single-cell RNA-seq
SSc breast: Breast Sensitivity Signature Collection
SP: Switch point
ST: Spatial transcriptomics
TC: Therapeutic cluster
TCGA: The Cancer Genome Atlas
TIS: Tumour inflammation signature
TLS: Tertiary lymphoid structure
TME: Tumour microenvironment
TNBC: Triple-negative breast cancer
UMAP: Uniform Manifold Approximation and Projection

## Declarations

### Ethics approval and consent to participate

Not applicable.

### Consent for publication

Not applicable.

### Availability of data and materials

We downloaded luminal and TNBC Visium ST data (CID4290, CID4535, 1142243F, 1160920F, CID4465 and CID44971) [5] from Zenodo (DOI: 10.5281/zenodo.4739739). HER2+ Visium ST data (738811QB, 1168993F and V19L29), licensed under the Creative Commons Attribution licence, were downloaded from the 10x Genomics Datasets portal [66–69]. All these data included raw counts and matching tissue images. Wu *et al.* dataset also included metadata information with pathologist annotations for each spot. 10x Genomics provided an image with pathologist annotations for patient 738811QB, and we manually labelled each spot using Loupe Browser v5.0.1 [111]. Additionally, we downloaded scRNA-seq data [5] from the Single Cell Portal (ID: SCP1039) to construct the spot deconvolution reference.

The SSc breast collection and Beyondcell objects generated in this work can be found on Zenodo (DOI: 10.5281/zenodo.10638905). The code used to create the SSc breast collection is available at https://github.com/cnio-bu/SSc-breast. The data preprocessing pipeline is accessible at https://github.com/cnio-bu/ST-preprocess. The code used to produce the final figures and tables is provided at https://github.com/cnio-bu/breast-bcspatial. Additionally, the Beyondcell and ggseabubble packages can be accessed at https://github.com/cniobu/beyondcell and https://github.com/mj-jimenez/ggseabubble.

### Authors’ contributions

MJJ-S designed the study, performed the analyses and wrote the manuscript under the supervision of GG-L and FA. SG-M computed the SSc breast signature collection and reviewed the manuscript. MR-F analysed patient V19L29 independently, providing insights that influenced the principal analysis. Beyondcell was updated and tested by MJJ-S and maintained by MJJ-S and SG-M. All authors read and approved the final manuscript.

## Acknowledgements

We thank all CNIO Bioinformatics Unit members for their support. We also thank Coral Fustero-Torre for her literature suggestions and fruitful discussions about the state of the art.

## References

1. Duffy MJ, Crown J. A personalized approach to cancer treatment: how biomarkers can help. Clin Chem. 2008;54:1770–9.

2. Burrell RA, McGranahan N, Bartek J, Swanton C. The causes and consequences of genetic heterogeneity in cancer evolution. Nature. 2013;501:338–45.

3. Wahida A, Buschhorn L, Fröhling S, Jost PJ, Schneeweiss A, Lichter P, et al. The coming decade in precision oncology: six riddles. Nat Rev Cancer. 2023;23:43–54.

4. Jiménez-Santos MJ, García-Martín S, Fustero-Torre C, Di Domenico T, Gómez-López G, Al-Shahrour F. Bioinformatics roadmap for therapy selection in cancer genomics. Mol Oncol. 2022;16:3881–908.

5. Wu SZ, Al-Eryani G, Roden DL, Junankar S, Harvey K, Andersson A, et al. A single-cell and spatially resolved atlas of human breast cancers. Nat Genet. 2021;53:1334–47.

6. Burguin A, Diorio C, Durocher F. Breast Cancer Treatments: Updates and New Challenges. J Pers Med. 2021;11:808.

7. Zagami P, Carey LA. Triple negative breast cancer: Pitfalls and progress. NPJ Breast Cancer. 2022;8:95.

8. Marusyk A, Janiszewska M, Polyak K. Intratumor Heterogeneity: The Rosetta Stone of Therapy Resistance. Cancer Cell. 2020;37:471–84.

9. Dagogo-Jack I, Shaw AT. Tumour heterogeneity and resistance to cancer therapies. Nat Rev Clin Oncol. 2018;15:81–94.

10. Dentro SC, Leshchiner I, Haase K, Tarabichi M, Wintersinger J, Deshwar AG, et al. Characterizing genetic intra-tumor heterogeneity across 2,658 human cancer genomes. Cell. 2021;184:2239–54.e39.

11. Barkley D, Moncada R, Pour M, Liberman DA, Dryg I, Werba G, et al. Cancer cell states recur across tumor types and form specific interactions with the tumor microenvironment. Nat Genet. 2022;54:1192–201.

12. Anderson NM, Simon MC. The tumor microenvironment. Curr Biol. 2020;30:R921–5.

13. de Visser KE, Joyce JA. The evolving tumor microenvironment: From cancer initiation to metastatic outgrowth. Cancer Cell. 2023;41:374–403.

14. Baghban R, Roshangar L, Jahanban-Esfahlan R, Seidi K, Ebrahimi-Kalan A, Jaymand M, et al. Tumor microenvironment complexity and therapeutic implications at a glance. Cell Commun Signal. 2020;18:59.

15. González-Silva L, Quevedo L, Varela I. Tumor Functional Heterogeneity Unraveled by scRNA-seq Technologies. Trends Cancer Res. 2020;6:13–9.

16. Seferbekova Z, Lomakin A, Yates LR, Gerstung M. Spatial biology of cancer evolution. Nat Rev Genet. 2023;24:295–313.

17. Williams CG, Lee HJ, Asatsuma T, Vento-Tormo R, Haque A. An introduction to spatial transcriptomics for biomedical research. Genome Med. 2022;14:68.

18. Fustero-Torre C, Jiménez-Santos MJ, García-Martín S, Carretero-Puche C, García-Jimeno L, Ivanchuk V, et al. Beyondcell: targeting cancer therapeutic heterogeneity in single-cell RNA-seq data. Genome Med. 2021;13:187.

19. Hao Y, Hao S, Andersen-Nissen E, Mauck WM 3rd, Zheng S, Butler A, et al. Integrated analysis of multimodal single-cell data. Cell. 2021;184:3573–87.e29.

20. Cable DM, Murray E, Zou LS, Goeva A, Macosko EZ, Chen F, et al. Robust decomposition of cell type mixtures in spatial transcriptomics. Nat Biotechnol. 2022;40:517–26.

21. Yoshihara K, Shahmoradgoli M, Martínez E, Vegesna R, Kim H, Torres-Garcia W, et al. Inferring tumour purity and stromal and immune cell admixture from expression data. Nat Commun. 2013;4:2612.

22. De Falco A, Caruso F, Su X-D, Iavarone A, Ceccarelli M. A variational algorithm to detect the clonal copy number substructure of tumors from scRNA-seq data. Nat Commun. 2023;14:1074.

23. Ritchie ME, Phipson B, Wu D, Hu Y, Law CW, Shi W, et al. limma powers differential expression analyses for RNA-sequencing and microarray studies. Nucleic Acids Res. 2015;43:e47.

24. Jang IS, Neto EC, Guinney J, Friend SH, Margolin AA. Systematic assessment of analytical methods for drug sensitivity prediction from cancer cell line data. Pac Symp Biocomput. 2014;63–74.

25. Basu A, Bodycombe NE, Cheah JH, Price EV, Liu K, Schaefer GI, et al. An interactive resource to identify cancer genetic and lineage dependencies targeted by small molecules. Cell. 2013;154:1151–61.

26. Seashore-Ludlow B, Rees MG, Cheah JH, Cokol M, Price EV, Coletti ME, et al. Harnessing Connectivity in a Large-Scale Small-Molecule Sensitivity Dataset. Cancer Discov. 2015;5:1210–23.

27. Rees MG, Seashore-Ludlow B, Cheah JH, Adams DJ, Price EV, Gill S, et al. Correlating chemical sensitivity and basal gene expression reveals mechanism of action. Nat Chem Biol. 2016;12:109–16.

28. Garnett MJ, Edelman EJ, Heidorn SJ, Greenman CD, Dastur A, Lau KW, et al. Systematic identification of genomic markers of drug sensitivity in cancer cells. Nature. 2012;483:570–5.

29. Yang W, Soares J, Greninger P, Edelman EJ, Lightfoot H, Forbes S, et al. Genomics of Drug Sensitivity in Cancer (GDSC): a resource for therapeutic biomarker discovery in cancer cells. Nucleic Acids Res. 2013;41:D955–61.

30. Iorio F, Knijnenburg TA, Vis DJ, Bignell GR, Menden MP, Schubert M, et al. A Landscape of Pharmacogenomic Interactions in Cancer. Cell. 2016;166:740–54.

31. Yu C, Mannan AM, Yvone GM, Ross KN, Zhang Y-L, Marton MA, et al. High-throughput identification of genotype-specific cancer vulnerabilities in mixtures of barcoded tumor cell lines. Nat Biotechnol. 2016;34:419–23.

32. Corsello SM, Nagari RT, Spangler RD, Rossen J, Kocak M, Bryan JG, et al. Discovering the anti-cancer potential of non-oncology drugs by systematic viability profiling. Nat Cancer. 2020;1:235–48.

33. Ghandi M, Huang FW, Jané-Valbuena J, Kryukov GV, Lo CC, McDonald ER 3rd, et al. Next-generation characterization of the Cancer Cell Line Encyclopedia. Nature. 2019;569:503–8.

34. Warren A, Chen Y, Jones A, Shibue T, Hahn WC, Boehm JS, et al. Global computational alignment of tumor and cell line transcriptional profiles. Nat Commun. 2021;12:22.

35. Larsson L, Franzén L, Ståhl PL, Lundeberg J. Semla: a versatile toolkit for spatially resolved transcriptomics analysis and visualization. Bioinformatics. 2023;39:btad626.

36. Korotkevich G, Sukhov V, Budin N, Shpak B, Artyomov MN, Sergushichev A. Fast gene set enrichment analysis. bioRxiv. 2021; doi:10.1101/060012.

37. Subramanian A, Tamayo P, Mootha VK, Mukherjee S, Ebert BL, Gillette MA, et al. Gene set enrichment analysis: a knowledge-based approach for interpreting genome-wide expression profiles. Proc Natl Acad Sci U S A. 2005;102:15545–50.

38. Liberzon A, Birger C, Thorvaldsdóttir H, Ghandi M, Mesirov JP, Tamayo P. The Molecular Signatures Database (MSigDB) hallmark gene set collection. Cell Syst. 2015;1:417–25.

39. Gillespie M, Jassal B, Stephan R, Milacic M, Rothfels K, Senff-Ribeiro A, et al. The reactome pathway knowledgebase 2022. Nucleic Acids Res. 2022;50:D687–92.

40. Yuan H, Yan M, Zhang G, Liu W, Deng C, Liao G, et al. CancerSEA: a cancer single-cell state atlas. Nucleic Acids Res. 2019;47:D900–8.

41. Schuetz CS, Bonin M, Clare SE, Nieselt K, Sotlar K, Walter M, et al. Progression-specific genes identified by expression profiling of matched ductal carcinomas in situ and invasive breast tumors, combining laser capture microdissection and oligonucleotide microarray analysis. Cancer Res. 2006;66:5278–86.

42. Buffa FM, Harris AL, West CM, Miller CJ. Large meta-analysis of multiple cancers reveals a common, compact and highly prognostic hypoxia metagene. Br J Cancer. 2010;102:428–35.

43. Gröger CJ, Grubinger M, Waldhör T, Vierlinger K, Mikulits W. Meta-analysis of gene expression signatures defining the epithelial to mesenchymal transition during cancer progression. PLoS One. 2012;7:e51136.

44. Ayers M, Lunceford J, Nebozhyn M, Murphy E, Loboda A, Kaufman DR, et al. IFN-γ-related mRNA profile predicts clinical response to PD-1 blockade. J Clin Invest. 2017;127:2930–40.

45. Schroth W, Büttner FA, Kandabarau S, Hoppe R, Fritz P, Kumbrink J, et al. Gene Expression Signatures of BRCAness and Tumor Inflammation Define Subgroups of Early-Stage Hormone Receptor-Positive Breast Cancer Patients. Clin Cancer Res. 2020;26:6523–34.

46. Cabrita R, Lauss M, Sanna A, Donia M, Skaarup Larsen M, Mitra S, et al. Tertiary lymphoid structures improve immunotherapy and survival in melanoma. Nature. 2020;577:561–5.

47. Wang W, Taufalele PV, Millet M, Homsy K, Smart K, Berestesky ED, et al. Matrix stiffness regulates tumor cell intravasation through expression and ESRP1-mediated alternative splicing of MENA. Cell Rep. 2023;42:112338.

48. Jin S, Plikus MV, Nie Q. CellChat for systematic analysis of cell-cell communication from single-cell and spatially resolved transcriptomics. bioRxiv. 2023; doi:10.1101/2023.11.05.565674.

49. R Core Team. R: A Language and Environment for Statistical Computing. Version 4. 2023. https://www.R-project.org/.

50. Kassambara A. rstatix: Pipe-Friendly Framework for Basic Statistical Tests. Version 0.7.2. 2023. https://CRAN.R-project.org/package=rstatix.

51. Wickham H, Averick M, Bryan J, Chang W, McGowan L, François R, et al. Welcome to the tidyverse. J Open Source Softw. 2019;4:1686.

52. Lawrence M, Huber W, Pagès H, Aboyoun P, Carlson M, Gentleman R, et al. Software for computing and annotating genomic ranges. PLoS Comput Biol. 2013;9:e1003118.

53. Kassambara A. ggpubr: “ggplot2” Based Publication Ready Plots. Version 0.6.0. 2023. https://CRAN.R-project.org/package=ggpubr.

54. Jimenez-Santos MJ. ggseabubble: R package for creating publication-ready bubble heatmaps. Version 1.0.0. 2023. https://github.com/mj-jimenez/ggseabubble.

55. Sjoberg D. ggsankey: Sankey, Alluvial and Sankey Bump Plots. Version 0.0.99999. 2023. https://github.com/davidsjoberg/ggsankey.

56. Larsson J. eulerr: Area-Proportional Euler and Venn Diagrams with Ellipses. Version 7.0.0. 2022. https://CRAN.R-project.org/package=eulerr.

57. Elosua-Bayes M, Nieto P, Mereu E, Gut I, Heyn H. SPOTlight: seeded NMF regression to deconvolute spatial transcriptomics spots with single-cell transcriptomes. Nucleic Acids Res. 2021;49:e50.

58. Gu Z, Eils R, Schlesner M. Complex heatmaps reveal patterns and correlations in multidimensional genomic data. Bioinformatics. 2016;32:2847–9.

59. Gu Z. Complex heatmap visualization. iMeta. 2022; doi:10.1002/imt2.43.

60. Gu Z, Gu L, Eils R, Schlesner M, Brors B. circlize Implements and enhances circular visualization in R. Bioinformatics. 2014;30:2811–2.

61. Wei T, Simko V. R package “corrplot”: Visualization of a Correlation Matrix. Version 0.92. 2021. https://github.com/taiyun/cor.

62. Neuwirth E. RColorBrewer: ColorBrewer Palettes. Version 1.1-3. 2022. https://CRAN.R-project.org/package=RColorBrewer.

63. Garnier S, Ross N, Rudis R, Camargo AP, Sciaini M, Scherer C. viridis(Lite) - Colorblind-Friendly Color Maps for R. Version 0.6.4. 2023. https://sjmgarnier.github.io/viridis/.

64. Pedersen TL. patchwork: The Composer of Plots. Version 1.1.3. 2023. https://CRAN.R-project.org/package=patchwork.

65. Johnston B. figpatch: Easily Arrange External Figures with Patchwork Alongside “ggplot2” Figures. Version 0.2. 2022. https://CRAN.R-project.org/package=figpatch.

66. 10x Genomics. Human Breast Cancer (Block A Section 1), Spatial Gene Expression Dataset by Space Ranger 1.1.0. 2020. https://www.10xgenomics.com/resources/datasets/human-breast-cancer-block-a-section-1-1-standard-1-1-0. Accessed 16 Jun 2023.

67. 10x Genomics. Human Breast Cancer (Block A Section 2), Spatial Gene Expression Dataset by Space Ranger 1.1.0. 2020. https://www.10xgenomics.com/resources/datasets/human-breast-cancer-block-a-section-2-1-standard-1-1-0. Accessed 16 Jun 2023.

68. 10x Genomics. Human Breast Cancer: Ductal Carcinoma In Situ, Invasive Carcinoma (FFPE), Spatial Gene Expression Dataset by Space Ranger 1.3.0. 2021. https://www.10xgenomics.com/resources/datasets/human-breast-cancer-ductal-carcinoma-in-situ-invasive-carcinoma-ffpe-1-standard-1-3-0. Accessed 16 Jun 2023.

69. 10x Genomics. Human Breast Cancer: Visium Fresh Frozen, Whole Transcriptome, Spatial Gene Expression Dataset by Space Ranger 1.3.0. 2022. https://www.10xgenomics.com/resources/datasets/human-breast-cancer-visium-fresh-frozen-whole-transcriptome-1-standard. Accessed 16 Jun 2023.

70. Marra A, Trapani D, Viale G, Criscitiello C, Curigliano G. Practical classification of triple-negative breast cancer: intratumoral heterogeneity, mechanisms of drug resistance, and novel therapies. NPJ Breast Cancer. 2020;6:54.

71. Helleday T, Petermann E, Lundin C, Hodgson B, Sharma RA. DNA repair pathways as targets for cancer therapy. Nat Rev Cancer. 2008;8:193–204.

72. Kao T-W, Bai G-H, Wang T-L, Shih I-M, Chuang C-M, Lo C-L, et al. Novel cancer treatment paradigm targeting hypoxia-induced factor in conjunction with current therapies to overcome resistance. J Exp Clin Cancer Res. 2023;42:171.

73. Foroutan M, Bhuva DD, Lyu R, Horan K, Cursons J, Davis MJ. Single sample scoring of molecular phenotypes. BMC Bioinformatics. 2018;19:404.

74. Bagaev A, Kotlov N, Nomie K, Svekolkin V, Gafurov A, Isaeva O, et al. Conserved pan-cancer microenvironment subtypes predict response to immunotherapy. Cancer Cell. 2021;39:845–65.e7.

75. Feng B, Wu J, Shen B, Jiang F, Feng J. Cancer-associated fibroblasts and resistance to anticancer therapies: status, mechanisms, and countermeasures. Cancer Cell Int. 2022;22:166.

76. Alshaker H, Thrower H, Pchejetski D. Sphingosine Kinase 1 in Breast Cancer-A New Molecular Marker and a Therapy Target. Front Oncol. 2020;10:289.

77. Beckwitt CH, Brufsky A, Oltvai ZN, Wells A. Statin drugs to reduce breast cancer recurrence and mortality. Breast Cancer Res. 2018;20:144.

78. Inasu M, Feldt M, Jernström H, Borgquist S, Harborg S. Statin use and patterns of breast cancer recurrence in the Malmö Diet and Cancer Study. Breast. 2022;61:123–8.

79. Damotte D, Warren S, Arrondeau J, Boudou-Rouquette P, Mansuet-Lupo A, Biton J, et al. The tumor inflammation signature (TIS) is associated with anti-PD-1 treatment benefit in the CERTIM pan-cancer cohort. J Transl Med. 2019;17:357.

80. Deng B, Zhao Z, Kong W, Han C, Shen X, Zhou C. Biological role of matrix stiffness in tumor growth and treatment. J Transl Med. 2022;20:1–15.

81. Levitin HM, Yuan J, Sims PA. Single-Cell Transcriptomic Analysis of Tumor Heterogeneity. Trends Cancer Res. 2018;4:264– 8.

82. Chen J, Wang X, Ma A, Wang Q-E, Liu B, Li L, et al. Deep transfer learning of cancer drug responses by integrating bulk and single-cell RNA-seq data. Nat Commun. 2022;13:6494.

83. Lotfollahi M, Klimovskaia Susmelj A, De Donno C, Hetzel L, Ji Y, Ibarra IL, et al. Predicting cellular responses to complex perturbations in high-throughput screens. Mol Syst Biol. 2023;19:e11517.

84. Pellecchia S, Viscido G, Franchini M, Gambardella G. Predicting drug response from single-cell expression profiles of tumours. BMC Med. 2023;21:476.

85. Giraldo NA, Sanchez-Salas R, Peske JD, Vano Y, Becht E, Petitprez F, et al. The clinical role of the TME in solid cancer. Br J Cancer. 2019;120:45–53.

86. Vitale I, Shema E, Loi S, Galluzzi L. Intratumoral heterogeneity in cancer progression and response to immunotherapy. Nat Med. 2021;27:212–24.

87. Tan K, Naylor MJ. Tumour Microenvironment-Immune Cell Interactions Influencing Breast Cancer Heterogeneity and Disease Progression. Front Oncol. 2022;12:876451.

88. Elhanani O, Ben-Uri R, Keren L. Spatial profiling technologies illuminate the tumor microenvironment. Cancer Cell. 2023;41:404–20.

89. Helmlinger G, Yuan F, Dellian M, Jain RK. Interstitial pH and pO2 gradients in solid tumors in vivo: high-resolution measurements reveal a lack of correlation. Nat Med. 1997;3:177–82.

90. Roussos ET, Condeelis JS, Patsialou A. Chemotaxis in cancer. Nat Rev Cancer. 2011;11:573–87.

91. Carmona-Fontaine C, Deforet M, Akkari L, Thompson CB, Joyce JA, Xavier JB. Metabolic origins of spatial organization in the tumor microenvironment. Proc Natl Acad Sci U S A. 2017;114:2934–9.

92. Kohli K, Pillarisetty VG, Kim TS. Key chemokines direct migration of immune cells in solid tumors. Cancer Gene Ther. 2022;29:10–21.

93. Ahmed MAM, Nagelkerke A. Current developments in modelling the tumour microenvironment in vitro: Incorporation of biochemical and physical gradients. Organs-on-a-Chip. 2021;3:100012.

94. Chen Z, Han F, Du Y, Shi H, Zhou W. Hypoxic microenvironment in cancer: molecular mechanisms and therapeutic interventions. Signal Transduct Target Ther. 2023;8:70.

95. Mahoney BP, Raghunand N, Baggett B, Gillies RJ. Tumor acidity, ion trapping and chemotherapeutics. I. Acid pH affects the distribution of chemotherapeutic agents in vitro. Biochem Pharmacol. 2003;66:1207–18.

96. Greijer AE, de Jong MC, Scheffer GL, Shvarts A, van Diest PJ, van der Wall E. Hypoxia-induced acidification causes mitoxantrone resistance not mediated by drug transporters in human breast cancer cells. Cell Oncol. 2005;27:43–9.

97. Rizzolio S, Giordano S, Corso S. The importance of being CAFs (in cancer resistance to targeted therapies). J Exp Clin Cancer Res. 2022;41:319.

98. Goyal Y, Busch GT, Pillai M, Li J, Boe RH, Grody EI, et al. Diverse clonal fates emerge upon drug treatment of homogeneous cancer cells. Nature. 2023;620:651–9.

99. Gatenby RA, Silva AS, Gillies RJ, Frieden BR. Adaptive therapy. Cancer Res. 2009;69:4894–903.

100. Piñeiro-Yáñez E, Jiménez-Santos MJ, Gómez-López G, Al-Shahrour F. In Silico Drug Prescription for Targeting Cancer Patient Heterogeneity and Prediction of Clinical Outcome. Cancers (Basel). 2019;11:1361.

101. Russell AJC, Weir JA, Nadaf NM, Shabet M, Kumar V, Kambhampati S, et al. Slide-tags enables single-nucleus barcoding for multimodal spatial genomics. Nature. 2024;625:101–9.

102. Nagendran M, Sapida J, Arthur J, Kamath G, Patterson D, Tentori A. Visium HD enables spatial discovery in FFPE human breast cancer at single-cell scale. 10x Genomics. 2023. https://pages.10xgenomics.com/rs/446-PBO-704/images/Monica_Visium%20HD%20final%20poster%20_SITC%20conference_2023.pdf?version=0. Accessed 10 Jan 2024.

103. Hu J, Coleman K, Zhang D, Lee EB, Kadara H, Wang L, et al. Deciphering tumor ecosystems at super resolution from spatial transcriptomics with TESLA. Cell Syst. 2023;14:404–17.e4.

104. Dries R, Zhu Q, Dong R, Eng C-HL, Li H, Liu K, et al. Giotto: a toolbox for integrative analysis and visualization of spatial expression data. Genome Biol. 2021;22:78.

105. Pham D, Tan X, Balderson B, Xu J, Grice LF, Yoon S, et al. Robust mapping of spatiotemporal trajectories and cell-cell interactions in healthy and diseased tissues. Nat Commun. 2023;14:7739.

106. Zhao E, Stone MR, Ren X, Guenthoer J, Smythe KS, Pulliam T, et al. Spatial transcriptomics at subspot resolution with BayesSpace. Nat Biotechnol. 2021;39:1375–84.

107. Vickovic S, Schapiro D, Carlberg K, Lötstedt B, Larsson L, Hildebrandt F, et al. Three-dimensional spatial transcriptomics uncovers cell type localizations in the human rheumatoid arthritis synovium. Commun Biol. 2022;5:129.

108. Long Y, Ang KS, Li M, Chong KLK, Sethi R, Zhong C, et al. Spatially informed clustering, integration, and deconvolution of spatial transcriptomics with GraphST. Nat Commun. 2023;14:1155.

109. Liu W, Liao X, Luo Z, Yang Y, Lau MC, Jiao Y, et al. Probabilistic embedding, clustering, and alignment for integrating spatial transcriptomics data with PRECAST. Nat Commun. 2023;14:296.

110. Hao Y, Stuart T, Kowalski MH, Choudhary S, Hoffman P, Hartman A, et al. Dictionary learning for integrative, multimodal and scalable single-cell analysis. Nat Biotechnol. 2023; doi:10.1038/s41587-023-01767-y.

111. 10x Genomics. Loupe Browser. Version 5.0.1. 2021. https://www.10xgenomics.com/products/visium-analysis#loupe-browser.

