## Supplementary Figures for "Spatial Transcriptomics in Breast Cancer Reveals Tumour Microenvironment-Driven Drug Responses and Clonal Therapeutic Heterogeneity"

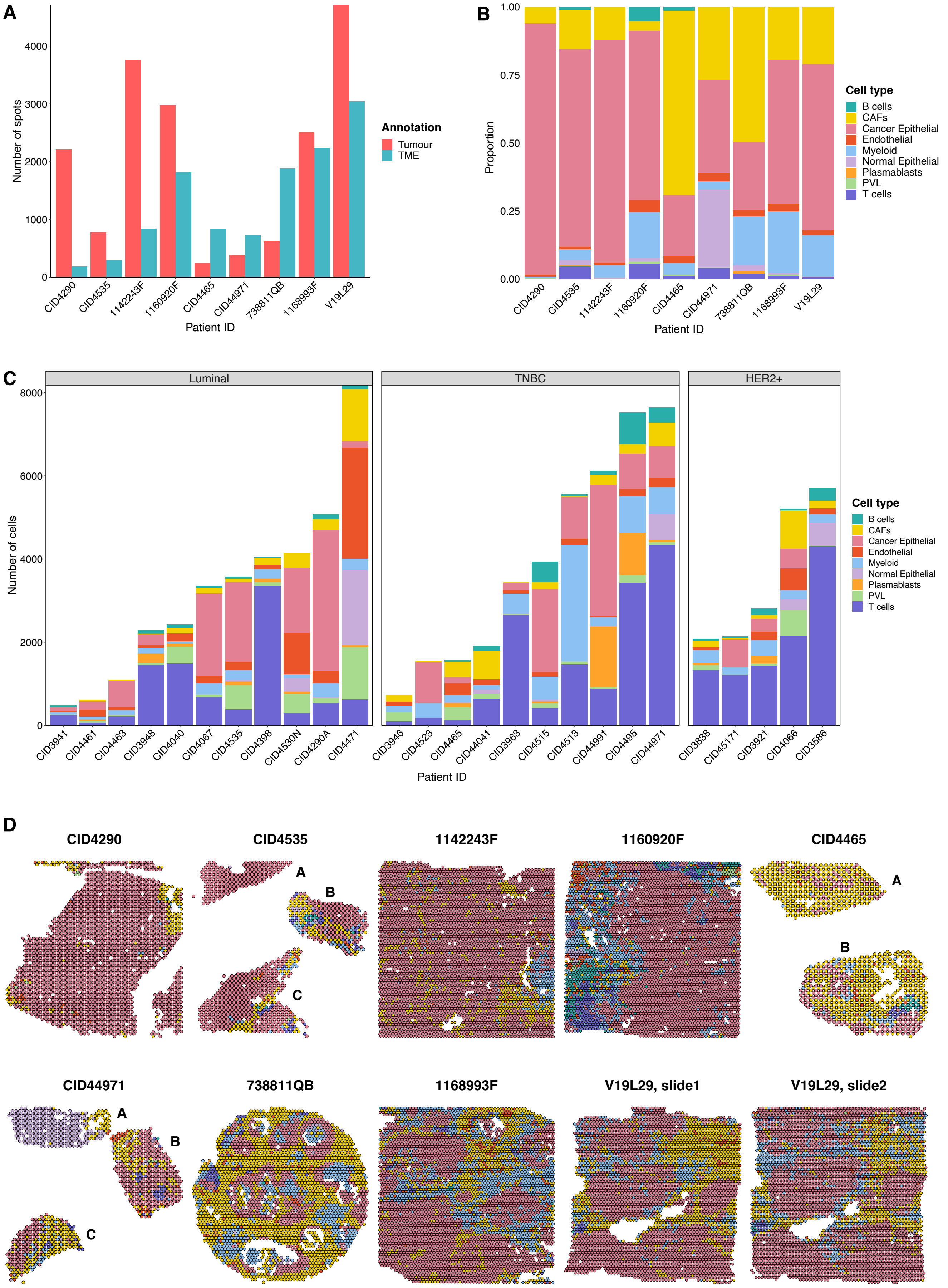

**Supplementary Figure S1. Cellular deconvolution of ST breast cancer samples. a)** Number of spots, coloured by tumour or TME annotation, in each ST sample. **b)** Proportion of deconvoluted cell types per ST sample. We labelled tumour spots as cancer cells and assigned TME spots to the non-malignant cell type with maximum deconvolved proportion. **c)** Number of cells, coloured by cell type, in each scRNA-seq sample used as a reference for spot deconvolution. **d)** Spatial projection of deconvoluted cell types in ST breast cancer samples. **ST:** Spatial transcriptomics; **TME:** Tumour microenvironment; **scRNA-seq:** single-cell RNA-seq; **CAFs:** Cancer-associated fibroblasts; **PVL**: Perivascular-like cells; **TNBC:** Triple-negative breast cancer; **HER2+:** Human epidermal growth factor receptor 2 positive.

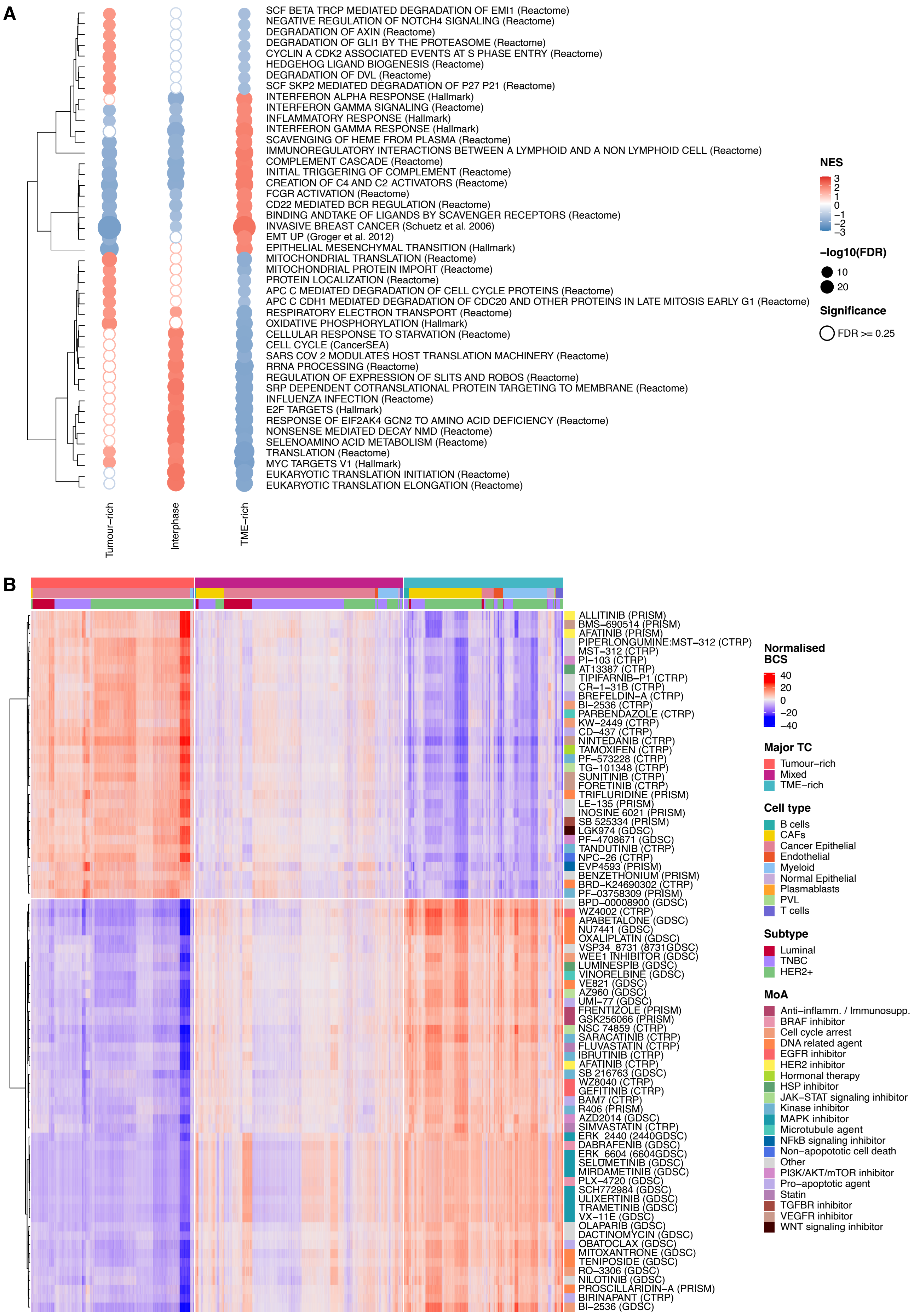

A
SCF BETA TRCP MEDIATED DEGRADATION OF EMI1 (Reactome)
NEGATIVE REGULATION OF NOTCH4 SIGNALING (Reactome)
DEGRADATION OF AXIN (Reactome)
DEGRADATION OF GLI1 BY THE PROTEASOME (Reactome)
CYCLIN A CDK2 ASSOCIATED EVENTS AT S PHASE ENTRY (Reactome)
HEDGEHOG LIGAND BIOGENESIS (Reactome)
DEGRADATION OF DVL (Reactome)
SCF SKP2 MEDIATED DEGRADATION OF P27 P21 (Reactome)
INTERFERON ALPHA RESPONSE (Hallmark)
INTERFERON GAMMA SIGNALING (Reactome)
INFLAMMATORY RESPONSE (Hallmark)
INTERFERON GAMMA RESPONSE (Hallmark)
SCAVENGING OF HEME FROM PLASMA (Reactome)
IMMUNOREGULATORY INTERACTIONS BETWEEN A LYMPHOID AND A NON LYMPHOID CELL (Reactome)
COMPLEMENT CASCADE (Reactome)
INITIAL TRIGGERING OF COMPLEMENT (Reactome)
CREATION OF C4 AND C2 ACTIVATORS (Reactome)
FCGR ACTIVATION (Reactome)
CD22 MEDIATED BCR REGULATION (Reactome)
BINDING ANDTAKE OF LIGANDS BY SCAVENGER RECEPTORS (Reactome)
INVASIVE BREAST CANCER (Schuetz et al. 2006)
EMT UP (Groger et al. 2012)
EPITHELIAL MESENCHYMAL TRANSITION (Hallmark)
MITOCHONDRIAL TRANSLATION (Reactome)
MITOCHONDRIAL PROTEIN IMPORT (Reactome)
PROTEIN LOCALIZATION (Reactome)
APC C MEDIATED DEGRADATION OF CELL CYCLE PROTEINS (Reactome)
APC C CDH1 MEDIATED DEGRADATION OF CDC20 AND OTHER PROTEINS IN LATE MITOSIS EARLY G1 (Reactome)
RESPIRATORY ELECTRON TRANSPORT (Reactome)
OXIDATIVE PHOSPHORYLATION (Hallmark)
CELLULAR RESPONSE TO STARVATION (Reactome)
CELL CYCLE (CancerSEA)
SARS COV 2 MODULATES HOST TRANSLATION MACHINERY (Reactome)
RRNA PROCESSING (Reactome)
REGULATION OF EXPRESSION OF SLITS AND ROBOS (Reactome)
SRP DEPENDENT COTRANSLATIONAL PROTEIN TARGETING TO MEMBRANE (Reactome)
INFLUENZA INFECTION (Reactome)
E2F TARGETS (Hallmark)
RESPONSE OF EIF2AK4 GCN2 TO AMINO ACID DEFICIENCY (Reactome)
NONSENSE MEDIATED DECAY NMD (Reactome)
SELENOAMINO ACID METABOLISM (Reactome)
TRANSLATION (Reactome)
MYC TARGETS V1 (Hallmark)
EUKARYOTIC TRANSLATION INITIATION (Reactome)
EUKARYOTIC TRANSLATION ELONGATION (Reactome)
Tumour–rich
Interphase
TME–rich
NES
3
2
1
0
−1
−2
−3
−log10(FDR)
10
20
Significance
FDR >= 0.25
B
ALLITINIB (PRISM)
BMS−690514 (PRISM)
AFATINIB (PRISM)
PIPERLONGUMINE:MST−312 (CTRP)
MST−312 (CTRP)
PI−103 (CTRP)
AT13387 (CTRP)
TIPIFARNIB−P1 (CTRP)
CR−1−31B (CTRP)
BREFELDIN−A (CTRP)
BI−2536 (CTRP)
PARBENDAZOLE (CTRP)
KW−2449 (CTRP)
CD−437 (CTRP)
NINTEDANIB (CTRP)
TAMOXIFEN (CTRP)
PF−573228 (CTRP)
TG−101348 (CTRP)
SUNITINIB (CTRP)
FORETINIB (CTRP)
TRIFLURIDINE (PRISM)
LE−135 (PRISM)
INOSINE 6021 (PRISM)
SB 525334 (PRISM)
LGK974 (GDSC)
PF−4708671 (GDSC)
TANDUTINIB (CTRP)
NPC−26 (CTRP)
EVP4593 (PRISM)
BENZETHONIUM (PRISM)
BRD−K24690302 (CTRP)
PF−03758309 (PRISM)
BPD−00008900 (GDSC)
WZ4002 (CTRP)
APABETALONE (GDSC)
NU7441 (GDSC)
OXALIPLATIN (GDSC)
VSP34 8731 (8731GDSC)
WEE1 INHIBITOR (GDSC)
LUMINESPIB (GDSC)
VINORELBINE (GDSC)
VE821 (GDSC)
AZ960 (GDSC)
UMI−77 (GDSC)
FRENTIZOLE (PRISM)
GSK256066 (PRISM)
NSC 74859 (CTRP)
SARACATINIB (CTRP)
FLUVASTATIN (CTRP)
IBRUTINIB (CTRP)
AFATINIB (CTRP)
SB 216763 (GDSC)
WZ8040 (CTRP)
GEFITINIB (CTRP)
BAM7 (CTRP)
R406 (PRISM)
AZD2014 (GDSC)
SIMVASTATIN (CTRP)
ERK 2440 (2440GDSC)
DABRAFENIB (GDSC)
ERK 6604 (6604GDSC)
SELUMETINIB (GDSC)
MIRDAMETINIB (GDSC)
PLX−4720 (GDSC)
SCH772984 (GDSC)
ULIXERTINIB (GDSC)
TRAMETINIB (GDSC)
VX−11E (GDSC)
OLAPARIB (GDSC)
DACTINOMYCIN (GDSC)
OBATOCLAX (GDSC)
MITOXANTRONE (GDSC)
TENIPOSIDE (GDSC)
RO−3306 (GDSC)
NILOTINIB (GDSC)
PROSCILLARIDIN−A (PRISM)
BIRINAPANT (CTRP)
BI−2536 (GDSC)
Normalised BCS
40
20
0
−20
−40
Major TC
Tumour−rich
Mixed
TME−rich
Cell type
B cells
CAFs
Cancer Epithelial
Endothelial
Myeloid
Normal Epithelial
Plasmablasts
PVL
T cells
Subtype
Luminal
TNBC
HER2+
MoA
Anti−inflamm. / Immunosupp.
BRAF inhibitor
Cell cycle arrest
DNA related agent
EGFR inhibitor
HER2 inhibitor
Hormonal therapy
HSP inhibitor
JAK−STAT signaling inhibitor
Kinase inhibitor
MAPK inhibitor
Microtubule agent
NFkB signaling inhibitor
Non−apopotoic cell death
Other
PI3K/AKT/mTOR inhibitor
Pro−apoptoic agent
Statin
TGFBR inhibitor
VEGFR inhibitor
WNT signaling inhibitor

**Supplementary Figure S2. Compartment-wise functional analysis and drug ranking. a)** Bubble heatmap depicting the 15 most significantly positively enriched pathways in each major TC identified by differential gene expression analysis and pre-ranked GSEA. Rows represent functional pathways, and columns represent the three major TCs. The colour of the bubble is proportional to the NES magnitude and the size of the bubble to the FDR-adjusted p-value. Empty bubbles represent non-significant results (FDR>=0.25). Rows are clustered according to the Euclidean distance between NES. **b)** Heatmap of drugs to which the tumour compartment displays specific sensitivity or insensitivity compared to the TME region. Drugs are clustered according to their BCS using Ward's method and labelled by MoA. Spots from all samples are ordered by major TC, cell type and breast cancer subtype. **TCs:** Therapeutic clusters; **GSEA:** Gene Set Enrichment Analysis; **NES:** Normalised Enrichment Score; **FDR:** False Discovery Rate; **TME:** Tumour microenvironment; **BCS:** Beyondcell Scores; **MoA:** Mechanism of action; **CAFs:** Cancer-associated fibroblasts; **PVL**: Perivascular-like cells; **TNBC:** Triple-negative breast cancer; **HER2+:** Human epidermal growth factor receptor 2 positive; **Anti-inflamm.:** Anti-inflammatory; **Immunosupp.:** Immunosuppressor.

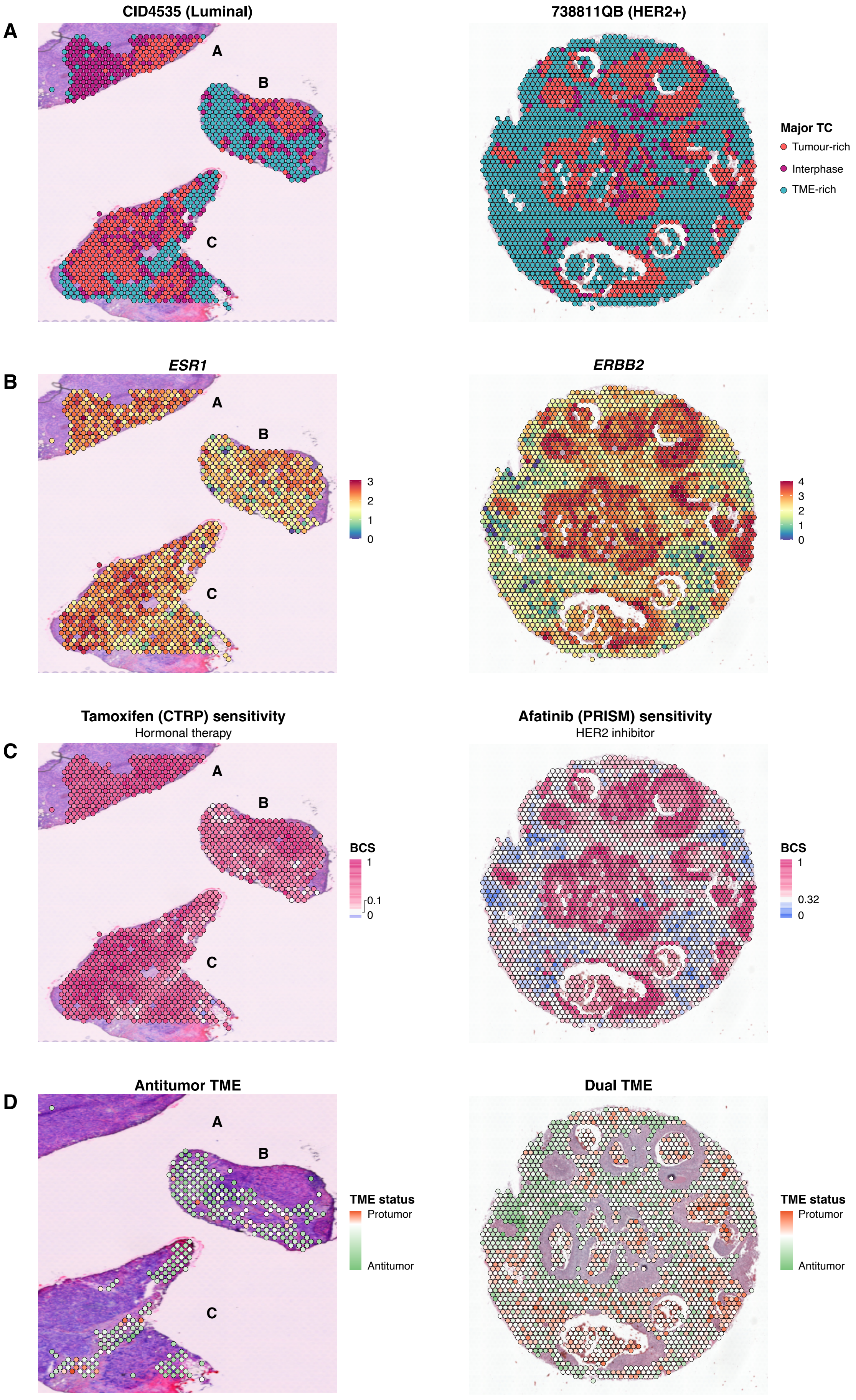

A
CID4535 (Luminal)
738811QB (HER2+)
Major TC
Tumour-rich
Interphase
TME-rich
B
ESR1
ERBB2
C
Tamoxifen (CTRP) sensitivity
Hormonal therapy
Afatinib (PRISM) sensitivity
HER2 inhibitor
BCS
1
0.1
0
0.32
D
Antitumor TME
Dual TME
TME status
Protumor
Antitumor

**Supplementary Figure S3. Individualised drug prescription in breast cancer subtypes.** Spatial projection of **a)** the 3 major TCs, **b)** the subtype-specific biomarker expression and **c)** the sensitivity to the standard-of-care treatment for luminal and HER2+ clinical subtypes. **d)** Spatial projection of the TME status of each TME spot. **TCs:** Therapeutic clusters; **HER2+:** Human epidermal growth factor receptor 2 positive; **TME:** Tumour microenvironment; ***ESR1*****:** Oestrogen Receptor gene; ***ERBB2*****:** Erb-B2 Receptor Tyrosine Kinase 2 gene; **CTRP:** Cancer Therapeutics Response Portal; **BCS:** Beyondcell Scores.

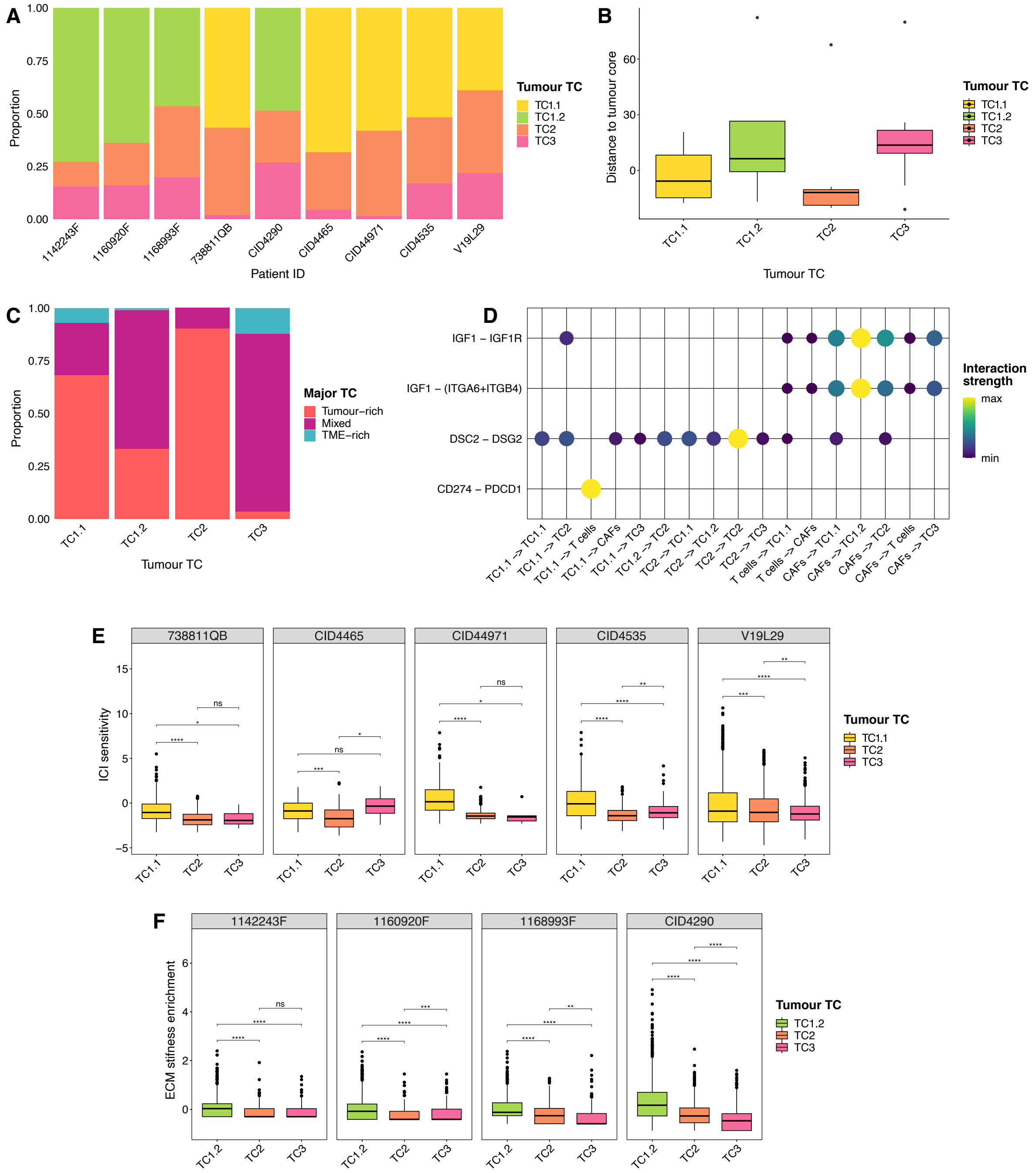

A
Proportion
1.00
0.75
0.50
0.25
0.00
1142243F
1160920F
1168993F
738811QB
CID4290
CID4465
CID44971
CID4535
V19L29
Patient ID
Tumour TC
TC1.1
TC1.2
TC2
TC3
B
Distance to tumour core
60
30
0
TC1.1
TC1.2
TC2
TC3
Tumour TC
C
Proportion
TC1.1
TC1.2
TC2
TC3
Tumour TC
Major TC
Tumour–rich
Mixed
TME–rich
D
IGF1 – IGF1R
IGF1 – (ITGA6+ITGB4)
DSC2 – DSG2
CD274 – PDCD1
TC1.1 -> TC1.1
TC1.1 -> TC2
TC1.1 -> T cells
TC1.1 -> CAFs
TC1.1 -> TC3
TC1.2 -> TC2
TC2 -> TC1.1
TC2 -> TC1.2
TC2 -> TC2
TC2 -> TC3
T cells -> TC1.1
T cells -> CAFs
CAFs -> TC1.1
CAFs -> TC1.2
CAFs -> TC2
CAFs -> T cells
CAFs -> TC3
Interaction strength
max
min
E
738811QB
CID4465
CID44971
CID4535
V19L29
ICI sensitivity
15
10
5
0
−5
ns
Tumour TC
TC1.1
TC2
TC3
F
1142243F
1160920F
1168993F
CID4290
ECM stifness enrichment
6
4
2
0
TC1.2
TC2
TC3
Tumour TC
TC1.2
TC2
TC3

**Supplementary Figure S4. Tumour TC characterisation. a)** Proportion of tumour TCs across all patients. **b)** Distribution of the region-wise mean radial distance to the tumour core of each tumour TC. To help visualisation, we plot the square root of mean radial distances multiplied by the original sign. **c)** Proportion of major TCs in each tumour TC. **d)** Bubble plot with significant (FDR<0.05) ligand-receptor pairs contributing to the signalling between tumour TCs, T cells and CAFs. The colour and size of the bubble are proportional to CellChat's interaction strength value, scaled for each ligand-receptor pair. **e)** Boxplots depicting a significantly higher ICI sensitivity in TC1.1 across patients. **f)** Boxplots depicting a significantly higher ECM stiffness in TC1.2 across patients. The box bounds the interquartile range divided by the median. Outliers are shown as dots. Differences in medians were tested with pairwise *post hoc* Wilcoxon rank-sum tests after significant Kruskal-Wallis tests. The p-values were adjusted using FDR correction for multiple testing (ns FDR>=0.05, *FDR<0.05, **FDR<0.01, ***FDR<0.001, ****FDR<0.0001). **TCs:** Therapeutic clusters; **FDR:** False Discovery Rate; **CAFs:** Cancer-associated fibroblasts; **ICI:** Immune checkpoint inhibitor; **ECM:** Extracellular matrix; **ns:** Not significant; **TME:** Tumour microenvironment.

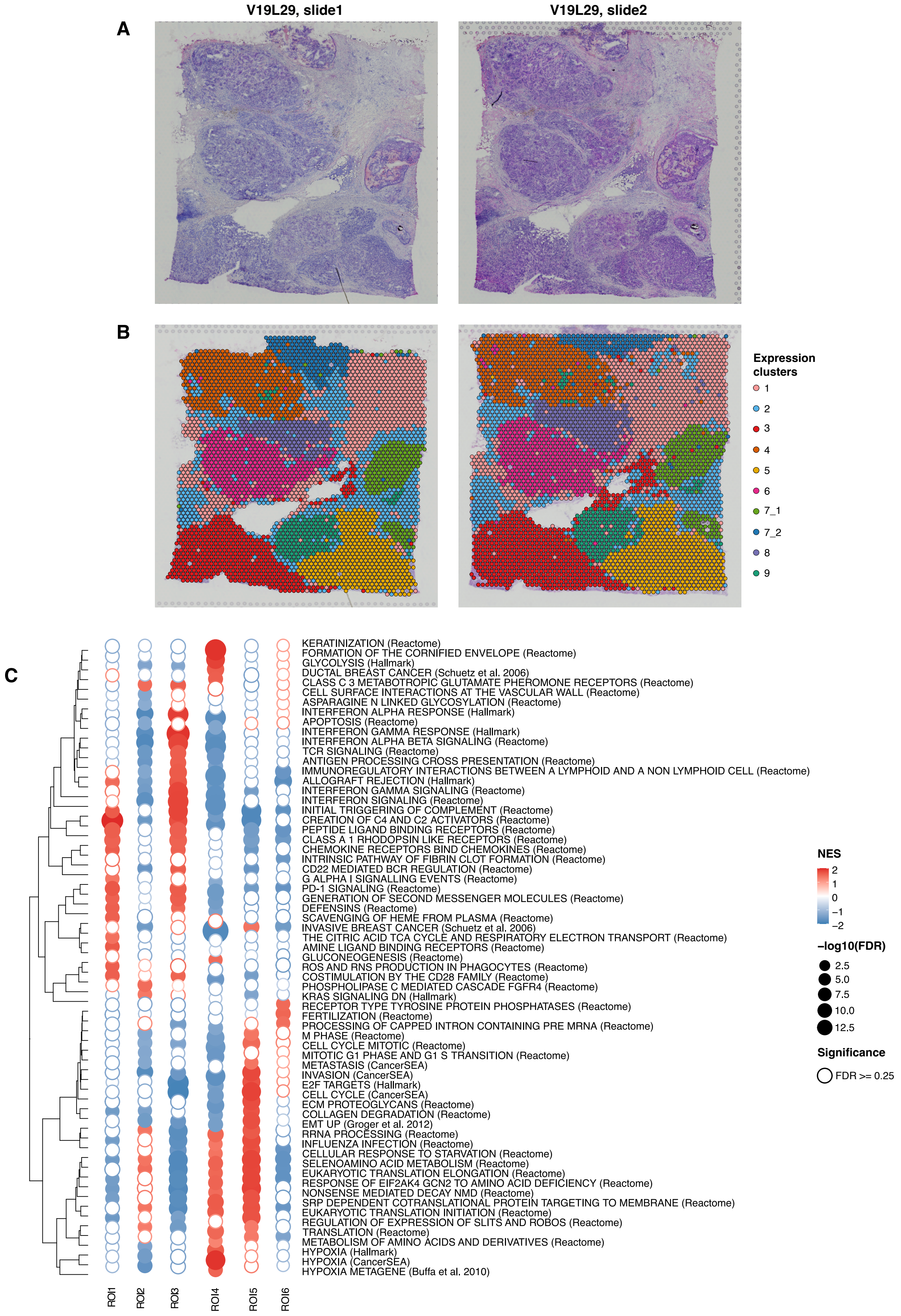
V19L29, slide1
V19L29, slide2
A
B
Expression clusters
1
2
3
4
5
6
7_1
7_2
8
9
C
KERATINIZATION (Reactome)
FORMATION OF THE CORNIFIED ENVELOPE (Reactome)
GLYCOLYSIS (Hallmark)
DUCTAL BREAST CANCER (Schuetz et al. 2006)
CLASS C 3 METABOTROPIC GLUTAMATE PHEROMONE RECEPTORS (Reactome)
CELL SURFACE INTERACTIONS AT THE VASCULAR WALL (Reactome)
ASPARAGINE N LINKED GLYCOSYLATION (Reactome)
INTERFERON ALPHA RESPONSE (Hallmark)
APOPTOSIS (Reactome)
INTERFERON GAMMA RESPONSE (Hallmark)
INTERFERON ALPHA BETA SIGNALING (Reactome)
TCR SIGNALING (Reactome)
ANTIGEN PROCESSING CROSS PRESENTATION (Reactome)
IMMUNOREGULATORY INTERACTIONS BETWEEN A LYMPHOID AND A NON LYMPHOID CELL (Reactome)
ALLOGRAFT REJECTION (Hallmark)
INTERFERON GAMMA SIGNALING (Reactome)
INTERFERON SIGNALING (Reactome)
INITIAL TRIGGERING OF COMPLEMENT (Reactome)
CREATION OF C4 AND C2 ACTIVATORS (Reactome)
PEPTIDE LIGAND BINDING RECEPTORS (Reactome)
CLASS A 1 RHODOPSIN LIKE RECEPTORS (Reactome)
CHEMOKINE RECEPTORS BIND CHEMOKINES (Reactome)
INTRINSIC PATHWAY OF FIBRIN CLOT FORMATION (Reactome)
CD22 MEDIATED BCR REGULATION (Reactome)
G ALPHA I SIGNALLING EVENTS (Reactome)
PD-1 SIGNALING (Reactome)
GENERATION OF SECOND MESSENGER MOLECULES (Reactome)
DEFENSINS (Reactome)
SCAVENGING OF HEME FROM PLASMA (Reactome)
INVASIVE BREAST CANCER (Schuetz et al. 2006)
THE CITRIC ACID TCA CYCLE AND RESPIRATORY ELECTRON TRANSPORT (Reactome)
AMINE LIGAND BINDING RECEPTORS (Reactome)
GLUCONEOGENESIS (Reactome)
ROS AND RNS PRODUCTION IN PHAGOCYTES (Reactome)
COSTIMULATION BY THE CD28 FAMILY (Reactome)
PHOSPHOLIPASE C MEDIATED CASCADE FGFR4 (Reactome)
KRAS SIGNALING DN (Hallmark)
RECEPTOR TYPE TYROSINE PROTEIN PHOSPHATASES (Reactome)
FERTILIZATION (Reactome)
PROCESSING OF CAPPED INTRON CONTAINING PRE MRNA (Reactome)
M PHASE (Reactome)
CELL CYCLE MITOTIC (Reactome)
MITOTIC G1 PHASE AND G1 S TRANSITION (Reactome)
METASTASIS (CancerSEA)
INVASION (CancerSEA)
E2F TARGETS (Hallmark)
CELL CYCLE (CancerSEA)
ECM PROTEOGLYCANS (Reactome)
COLLAGEN DEGRADATION (Reactome)
EMT UP (Groger et al. 2012)
RRNA PROCESSING (Reactome)
INFLUENZA INFECTION (Reactome)
CELLULAR RESPONSE TO STARVATION (Reactome)
SELENOAMINO ACID METABOLISM (Reactome)
EUKARYOTIC TRANSLATION ELONGATION (Reactome)
RESPONSE OF EIF2AK4 GCN2 TO AMINO ACID DEFICIENCY (Reactome)
NONSENSE MEDIATED DECAY NMD (Reactome)
SRP DEPENDENT COTRANSLATIONAL PROTEIN TARGETING TO MEMBRANE (Reactome)
EUKARYOTIC TRANSLATION INITIATION (Reactome)
REGULATION OF EXPRESSION OF SLITS AND ROBOS (Reactome)
TRANSLATION (Reactome)
METABOLISM OF AMINO ACIDS AND DERIVATIVES (Reactome)
HYPOXIA (Hallmark)
HYPOXIA (CancerSEA)
HYPOXIA METAGENE (Buffa et al. 2010)
ROI1
ROI2
ROI3
ROI4
ROI5
ROI6
NES
2
1
0
−1
−2
−log10(FDR)
2.5
5.0
7.5
10.0
12.5
Significance
FDR >= 0.25

**Supplementary Figure S5. Definition and biological characterisation of patient V19L29 tumoural ROIs. a)** H&E staining of patient V19L29 slides showing several well-delimited globular ducts. **b)** Spatial projection of expression clusters. **c)** Bubble heatmap depicting the 15 most significantly positively enriched pathways in each tumoural ROI, as identified by differential gene expression analysis and pre-ranked GSEA. Rows represent functional pathways, and columns represent the ROIs. The colour of the bubble is proportional to the NES magnitude and the size of the bubble to the FDR-adjusted p-value. Empty bubbles represent non-significant results (FDR>=0.25). Rows are clustered according to the Euclidean distance between NES. **ROI:** Region of interest; **H&E:** Hematoxylin-Eosin; **GSEA:** Gene Set Enrichment Analysis; **NES:** Normalised Enrichment Score; **FDR:** False Discovery Rate.
